# Coexistence of fast and slow gamma oscillations in one population of inhibitory spiking neurons

**DOI:** 10.1101/687624

**Authors:** Hongjie Bi, Marco Segneri, Matteo di Volo, Alessandro Torcini

## Abstract

Oscillations are a hallmark of neural population activity in various brain regions with a spectrum covering a wide range of frequencies. Within this spectrum gamma oscillations have received particular attention due to their ubiquitous nature and to their correlation with higher brain functions. Recently, it has been reported that gamma oscillations in the hippocampus of behaving rodents are segregated in two distinct frequency bands: slow and fast. These two gamma rhythms correspond to different states of the network, but their origin has been not yet clarified. Here, we show theoretically and numerically that a single inhibitory population can give rise to coexisting slow and fast gamma rhythms corresponding to collective oscillations of a balanced spiking network. The slow and fast gamma rhythms are generated via two different mechanisms: the fast one being driven by the coordinated tonic neural firing and the slow one by endogenous fluctuations due to irregular neural activity. We show that almost instantaneous stimulations can switch the collective gamma oscillations from slow to fast and vice versa. Furthermore, to make a closer contact with the experimental observations, we consider the modulation of the gamma rhythms induced by a slower (theta) rhythm driving the network dynamics. In this context, depending on the strength of the forcing and the noise amplitude, we observe phase-amplitude and phase-phase coupling between the fast and slow gamma oscillations and the theta forcing. Phase-phase coupling reveals on average different theta-phases preferences for the two coexisting gamma rhythms joined to a wide cycle-to-cycle variability.

## I. INTRODUCTION

The emergence of collective oscillations in complex system has been a subject largely studied in the last decades from an experimental as well as from a theoretical point of view, for a recent review see [1]. In particular, the transition from asynchronous to collective dynamics in networks of heterogeneous oscillators has been characterized in terms of methods borrowed from statistical mechanics [2–4] and nonlinear dynamics [5–7]. Exact analytic techniques to reduce the infinite dimensional dynamics of globally coupled inhomogeneous phase oscillators to few mean field variables have became available in the last decade [8] allowing for noticeable progresses in the field [1]. Quite recently, these reduction techniques have been applied to globally coupled spiking neural networks [9], thus opening new perspectives for the study of large ensembles of spiking neurons and for the understanding of the mechanisms underlying brain rhythms.

Oscillatory dynamics is fundamental for the functioning of the mammalian brains, rhythms ranging from 1 to 500 Hz have been measured at a mesoscopic level, corresponding to the dynamics of neural populations, by employing electroencephalography (EEG), magnetoencephalography (MEG), or local field potential (LFP) [10].

In particular, gamma oscillations (30-100 Hz) have been suggested to underlie various cognitive and motor functions. Oscillations in the gamma band have been related to attention selection [11], memory formation and retrieval [12, 13], binding mechanisms for sensory awareness [14], and human focal seizures [15].

Gamma oscillations have been observed in many areas of the brain and their emergence has been shown to be crucially dependent on inhibitory networks [16, 17]. By following [16] gamma oscillations in purely inhibitory networks can emerge only via two mechanisms: the single neurons can fire periodically locked in phase [18] or each neuron can have irregular activity, but sufficiently strong recurrent interactions can render the asynchronous state unstable against fluctuations and collective oscillations (COs) can arise [19–21]. On one hand, the role of the synaptic mechanisms in promoting tonic synchronization in the gamma range has been clarified in [17, 22]. On the other hand, fast network oscillations with irregular neural discharges can emerge when the neurons are operating in the so-called balanced state [23–27]. A typical cortical state, where the balance of excitation and inhibition allows for a healthy activity of the brain. The balanced state has been observed *in vitro* and *in vivo* experiments in the cerebral cortex [28, 29] and reported in simulations of networks of excitatory and inhibitory spiking neurons [20, 30, 31] as well as of purely inhibitory circuits driven by external excitatory currents [32, 33].

Gamma oscillations are usually modulated by theta oscillations in the hippocampus during locomotory actions and rapid eye movement (REM) sleep, theta frequencies correspond to 4-12 Hz in rodents [34, 35] and to 1-4 Hz in humans [36, 37]. Two mechanisms of entrainment (or cross-frequency coupling) between theta and gamma oscillations have been reported : namely, phase-amplitude (P-A) and phase-phase (P-P) coupling. The P-A coupling (or theta-nested gamma oscillations) corresponds to the fact that the phase of the theta-oscillation modifies the amplitude of the gamma waves [38, 39], while P-P coupling refers to n:m phase locking between gamma and theta phase oscillations [35, 40].

Recently, the co-existence of gamma oscillations in three distinct bands has been reported for the cornu ammonis area 1 (CA1) of the hippocampus [35]: namely, a slow one (≃ 30-50 Hz), a fast (or intermediate) one (≃ 50-90 Hz), and a so called *ε*-band (≃ 90-150 Hz). However, only the two lower bands show a clear correlation (P-P coupling) with the theta rhythm during maze exploration and REM sleep, thus suggesting their functional relevance [35]. There are several further evidences that these two gamma bands correspond to different states of the hippocampal network [41]. In particular, in freely behaving rats place cells code differently the space location and the running speed during theta-nested slow or fast gamma rhythms [41–43]. Moreover, gamma rhythms with similar low and high frequencies subtypes occur in many other brain regions, besides the hippocampus [44, 45]. Despite their relevance, the mechanisms behind the emergence of these two distinct gamma bands are not yet clarified.

For what concerns the hippocampus, experiments show that slow gamma rhythms couple the activity of the CA1 area to synaptic inputs from CA3, while fast gamma rhythms in CA1 are entrained by inputs from medial Entorinhal Cortex (mEC) [45]. Slow and fast oscillations have been recorded also in CA3, where fast gamma are entrained by synaptic inputs from mEC [46]. These findings suggest that CA3-activated interneurons drive slow gamma, while mEC-activated interneurons drive fast gamma. Nonetheless, it has been shown that a substantial proportion of CA1 interneurons phase-lock to both slow and fast gamma LFP oscillations [35, 46, 47]. Therefore, as suggested by L.L. Colgin in [45], such interneurons may be part of a network that can generate either slow or fast gamma, depending on the state of the network. Furthermore there are experimental evidences that gamma rhythms can be generated locally *in vitro* in the CA1, as well as in the CA3 and mEC, thanks to optogenetic stimulations [39, 48, 49] or pharmacological manipulations, but at lower gamma frequencies with respect to optogenetics [50–53]. A recent theoretical work has analysed the emergence of gamma oscillations in a neural circuit composed by two populations of interneurons with fast and slow synaptic time scales [54]. Based on the results of this idealized rate model and on the analysis of experimental data sets for the CA1 area the authors showed that multiple gamma bands can arise locally without being the reflection of feedfoward inputs.

In the present work, we show, for the first time to our knowledge, that a single inhibitory population, characterized by only one synaptic time, can display coexisting fast and slow gamma COs corresponding to different network states. In particular, the slow gamma oscillations are associated to irregular spiking behaviours and fluctuations driven, while the fast gamma oscillations coexist with a much more regular neural dynamics and they can be characterized as mean driven [55, 56]. Furthermore, in presence of theta forcing we observe different theta-gamma cross-frequency coupling scenarios depending on the forcing amplitude. For small amplitudes we have theta-nested gamma oscillations resembling those reported for various brain areas *in vitro* under optogenetic sinusoidal theta-stimulation [39, 48, 49]. At larger amplitudes the two types of gamma COs phase lock to the theta rhythm, similarly to what has been reported experimentally for the CA1 region of the hippocampus [35, 46]. More specifically we have studied balanced sparse inhibitory networks of quadratic integrate-and-fire (QIF) neurons pulse coupled via inhibitory post-synaptic potentials (IPSPs), characterized by a finite synaptic time scale. For this sparse network we derived an effective mean-field (MF) by employing recently developed reduction techniques for QIF networks [9, 21, 57, 58]. In the MF model, in proximity of the sub-critical Hopf bifurcations, we report regions of bistability involving one stable focus and one stable limit cycle. In direct simulations of the corresponding spiking network we observe the coexistence of two distinct COs with frequencies in the slow and fast gamma band. The slow gamma COs are due to the microscopic irregular dynamics, characteristic of the balanced dynamics, which turns the damped oscillations towards the MF focus in sustained COs. The fast gamma COs are instead related to the oscillatory branch emerging via the sub-critical Hopf bifurcation from the asynchronous state. The network can be driven from one kind of COs to the other by transiently stimulating the neurons. In presence of a theta forcing nested gamma oscillations characterized by a P-A coupling appear for small forcing amplitudes, while at intermediate amplitudes slow and fast gamma phases lock to the theta phase displaying P-P coupling between the rhythms. For even larger amplitudes only fast gamma are observables with a maximal power in correspondence of the maximum of the stimulation.

The paper is organized as follows. In Sec. II, we introduce the model for an inhibitory sparse balanced network of QIF neurons as well as the macroscopic and microscopic indicators employed to characterize its dynamics. Section III is devoted to the derivation of the corresponding effective MF model and to the linear stability analysis of the asynchronous state. Simulation results for the network for high and low structural heterogeneity are reported in Section IV and compared with MF forecasts. The coexistence and transitions from slow (fast) to fast (slow) gamma oscillations is analyzed in Section V together with the cross-frequency coupling between theta and gamma oscillations. A concise discussion of the results and of possible future developments is reported in Section VI. Finally, Appendix A is devoted to the analysis of coexisting gamma oscillations in Erdös-Reniy networks, while Appendix B discusses of a general mechanism for the coexistence of noise-driven and tonic oscillations.

## II. METHODS

### A. The network model

We consider *N* inhibitory pulse-coupled QIF neurons [59] arranged in a random sparse balanced network. The membrane potential of each neuron evolves according to the following equations:

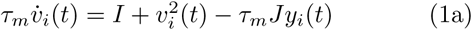

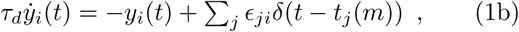

where *τ*_*m*_ = 15 ms represents the membrane time constant, *I* an external DC current, encompassing the effect of distal excitatory inputs and of the internal neural excitability. The last term in (1a) is the inhibitory synaptic current, with *J* being the synaptic coupling and *y*_*i*_ the synaptic field seen by neuron *i*. Whenever the membrane potential *v*_*i*_ reaches infinity a spike is emitted and *v*_*i*_ resetted to −∞.

The field *y*_*i*_ is the linear super-position of all the exponential IPSPs *s*(*t*) = exp (−*t/τ*_*d*_) received by the neuron *i* from its pre-synaptic neurons in the past, namely

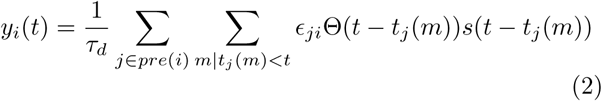

where *τ*_*d*_ is the synaptic time constant, *t*_*j*_(*m*) the spike time of the *m*-th spike delivered by the *j*-th neuron, Θ(*t*) is the Heaviside function and *ϵ*_*ji*_ is the adjacency matrix of the network. In particular, *ϵ*_*ji*_ = 1 (0) if a connection from node *j* to *i* exists (or not) and *k*_*i*_ = ∑ *ϵ*_*ji*_ is the number of pre-synaptic neurons connected to neuron *i*, or in other terms its in-degree.

In order to compare the simulation results with an exact MF recently derived [9, 21, 57], we consider sparse networks where the in-degrees *k*_*i*_ are extracted from a Lorentzian distribution

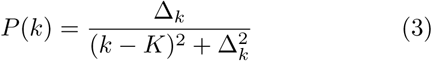

peaked at *K* and with a half-width half-maximum(HWHM) Δ_*k*_, the parameter Δ_*k*_ measures the level of structural heterogeneity in the network, and analogously to Erdös-Renyi networks we ass<u>u</u>med the following scaling for the HWHM 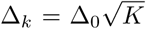. The DC current and the synaptic couplin<u>g</u> are rescaled wit<u>h</u> the median in degree *K* as 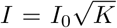 and 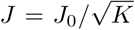, as usually done to achieve a self-sustained balanced state for sufficiently large in degrees [23–25, 27, 31]. In this paper we will usually consider *I*_0_ = 0.25, *N* = 10, 000 and *K* = 1, 000, unless stated otherwise.

### B. Simulation Protocols

The network dynamics is integrated by employing a standard Euler scheme with an integration time step Δ*t* = *τ*_*m*_*/*10000. The coexistence of solutions in proximity of a sub-critical Hopf bifurcation is analysed by performing adiabatic network simulations where a control parameter (e.g. the synaptic time *τ*_*d*_) is slowly varied. In particular, these are performed by starting with a initial value of 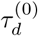 and arriving to a final value 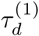 in *M* steps, each time increasing *τ*_*d*_ by 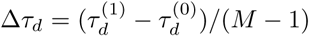. Once the final value 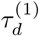 is reached, the synaptic time is decreased in steps Δ*τ*_*d*_ down to 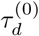. Each step corresponds to a simulation for a time *T*_*s*_ = 90 s during which the quantities of interest are measured, after discarding a transient *T*_*t*_ = 15 s. The initial condition for the system at each step is its final configuration at the previous step.

For what concerns the analysis of the crossing times *t*_*c*_ from slow (fast) to fast (slow) gamma in a bistable regime, reported in Section V A, we proceeded as follows. Let us first consider the transition from slow to fast gamma COs. We initialize the system in the slow gamma state at a current *I*_0_ ≡ *I*_1_ ensuring the bistability of the dynamics. Then we increase the DC current to a value *I*_0_ ≡ *I*_2_ for a time interval *T*_*P*_, after that time we return to the original value *I*_0_ ≡ *I*_1_ and we check, after a period of 1.5 s, if the system is in the slow or fast gamma regime. Then we repeat the process *M* = 30 times for each considered value of *T*_*P*_ and we measure the corresponding transition probability. The crossing time *t*_*c*_ is defined as the minimal *T*_*P*_ giving 80 % of probability that the transition will take place. To analyse the transition from fast to slow, we initialize the system in the fast gamma state at a DC current *I*_1_, we decrease the current to a value *I*_0_ ≡ *I*_3_ for time *T*_*P*_ and the we proceed as before. To examine the influence of noise on such transitions we added to the membrane potential evolution a noise term of zero average and amplitude *A*_*n*_.

### C. Indicators

To characterize the collective dynamics in the network we measure the mean membrane potential 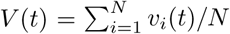, the instantaneous firing rate *R*(*t*), corresponding to the number of spikes emitted per unit of time and per neuron, as well as the mean synaptic field 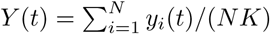 [60].

The microscopic activity can be analyzed by considering the inter-spike interval (ISI) distribution as characterized by the coefficient of variation *cv*_*i*_ for each neuron *i*, which is the ratio between the standard deviation and the mean of the ISIs associated to the train of spikes emitted by the considered neuron. In particular, we will characterize each network in terms of the average coefficient of variation defined as *CV* = ∑_*i*_ *cv*_*i*_ */N*. Time averages and fluctuations are usually estimated on time intervals *T*_*s*_ ≃ 90 s, after discarding transients *T*_*t*_ ≃ 15 s.

Phase entrainement between an external forcing characterized by its phase *θ*(*t*) and the collective oscillations induced in the network can be examined by considering the following phase difference:

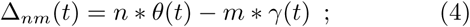

where *γ*(*t*) is the phase of the COs defined by considering the time occurrences *T*_*k*_ of the *k* maximum of the instantaneous firing rate *R*(*t*) of the network, namely *γ*(*t*) = 2*π*(*t* − *T*_*k*_)*/*(*T*_*k*+1_ − *T*_*k*_) with *t* ∈ [*T*_*k*_, *T*_*k*+1_] [61]. We have a *n* : *m* phase locking whenever the phase difference (4) is bounded during the time evolution, i.e. | Δ_*nm*_(*t*)| < *const*.

This somehow qualitative criterion can be made more quantitative by considering statistical indicators measuring the level of *n* : *m* synchronization for irregular/noisy data. In particular, an indicator based on the Shannon entropy has been introduced in [40], namely

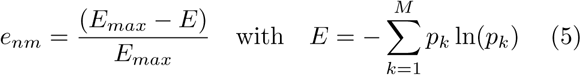

where *E* is the entropy associated to the distribution of Δ_*nm*_(*t*) and *E*_*max*_ = ln(*M*) with *M* number of bins.

The degree of synchronization among the phases can be also measured by the so-called Kuramoto order parameter, namely [35, 62]

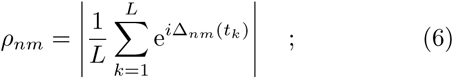

where | ·| represents the modulus and 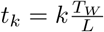 are *L* successive equispaced times within the considered time window *T*_*W*_. For completely desynchronized phases 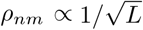, while partial (full) synchronization will be observable whenever *ρ*_*nm*_ is finite (one).

To assess the stationarity and the statistical significance of the obtained data we measured the above indicators within a time window *T*_*W*_ and we averaged the results over several distinct time windows in order to obtain also the corresponding error bars. Furthermore, to avoid the detection of spurious phase locking due to noise or band-pass filtering one should derive significance levels 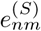 and 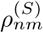 for each *n* : *m* phase locking indicators *e*_*nm*_ and *ρ*_*nm*_ [40, 63]. The significance levels have beeen estimated by considering surrogate data obtained by randomly shuffling the original time stamps of one of the two considered phases. Moreover, by following [63] we considered also other two types of surrogates for the generation of Δ_*nm*_(*t*) (4) within a certain time window *T*_*W*_. These are the time-shift surrogate, obtained by time shifting the origin of one time series for the phases with respect to the original one in the definiton of (4) and the random permutation surrogate, obtained by randomly choosing the origins of two time windows of duration *T*_*W*_ to estimate Δ_*nm*_(*t*).

## III. EFFECTIVE MEAN-FIELD MODEL FOR A SPARSE QIF NETWORK

By following [21] we derive an effective MF formulation for the model (1). As a starting point we consider an exact macroscopic model recently derived for fully coupled networks of pulse-coupled QIF [9], in particular we focus on inhibitory neurons coupled via exponentially decaying IPSPs [57]. For a structurally inhomogeneous network made of identical QIF neurons, with the synaptic couplings randomly distributed according to a Lorentzian, the MF dynamics can be expressed in terms of only three collective variables (namely, *V*, *R* and *Y*), as follows :

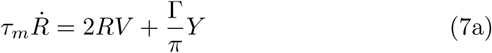

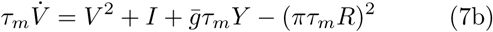

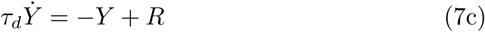

where 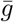 is the median and Γ the HWHM Lorentzian distribution of the synaptic couplings.

At a mean-field level, the above formulation can be applied to a sparse network, indeed the quenched disorder in the connectivity distribution can be rephrased in terms of a random synaptic coupling. Namely, each neuron *i* is subject in average to an inhibitory synaptic current of amplitude 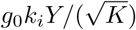 proportional to its indegree *k*_*i*_. Therefore at a first level of approximation we can consider the neurons as fully coupled, but with random values of the coupling distributed as a Lorentzian of median 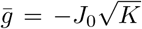 and HWHM Γ = *J*_0_Δ_0_. The MF formulation (7) takes now the expression:

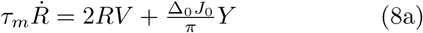

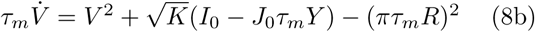

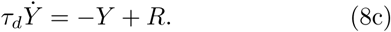

As verified in [21] for instantaneous PSPs this formulation represents a quite good guidance for the understanding of the emergence of sustained COs in the network, despite the fact that the MF asymptotic solutions are always stable foci. Instead in the present case, analogously to what found for structurally homogeneous networks of heterogeneous neurons in [57], we observe that for IPSPs of finite duration oscillations can emerge in the network as well as in the mean-field, as shown in Fig. 1. The data reported in the figure confirm that the MF formulation (8), despite not including current fluctuations, reproduces quite well the macroscopic evolution of the network in the oscillatory regime also for a sparse network.

**FIG. 1.**
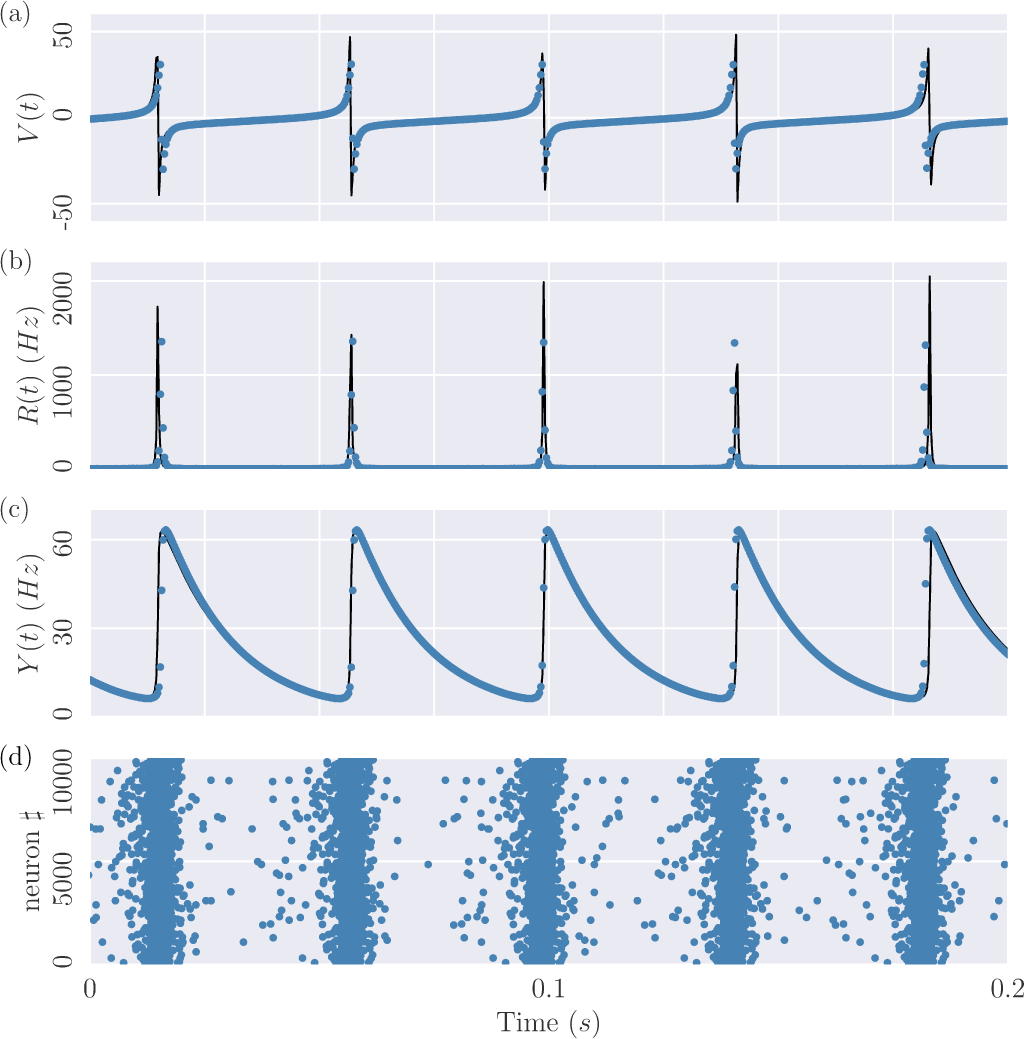
Comparison of the spiking dynamics with the mean-field results. Collective variables *V* (a), *R* (b) and *Y* (c) versus time, obtained from simulations of the spiking network (1) (blue circles) as well as from the MF formulation (8) (black line). In (d) the corresponding raster plot is also displayed, revealing clear COs with frequency *ν*_*OSC*_ ∼ 24 Hz. Dynamics of the network of *N* = 10000 neurons with median in-degree *K* = 1000 and Δ_0_ = 0.3. Other parameters are *I*_0_ = 0.25, *J*_0_ = 1.0 and *τ*_*d*_ = 15 ms.

Therefore we can safely employ such effective MF model to interpret the phenomena observed in the spiking network and to obtain theoretical predictions for its dynamics.

In the next two subsections we will firstly study analytically the linear stability of the asynchronous state, which corresponds to a fixed point of (8), and then we will describe the bifurcation and phase diagrams associated to the MF model (8).

### A. Linear stability of the asynchronous state

The fixed point solution (*V* *, *R*\*, *Y* *) of (8) is given by:

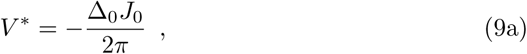

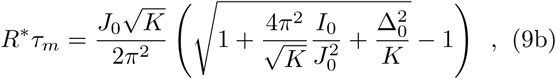

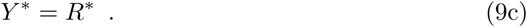

By performing a linear stability analysis around the fixed point solution (*V* *, *R*\*, *Y* *) we obtain the following secular equation:

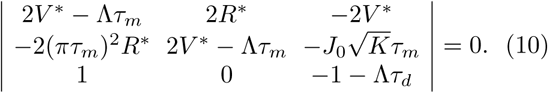

in a more explicit form this is

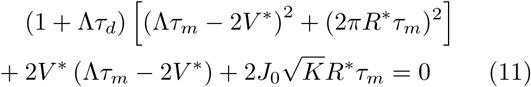

In the present case, for inhibitory coupling (i.e. *J*_0_ > 0) the solutions of the cubic equation (11) are one real and two complex conjugates. The real one is always negative therefore irrelevant for the stability analysis, while the couple of complex eigenvalues Λ = Λ_*R*_ ± *i*Λ_*I*_ can cross the imaginary axes giving rise to oscillatory behaviours via Hopf bifurcations. The presence of the two complex conjugate eigenvalues implies that whenever the asynchronous state is stable, this is always a focus characterized by a frequency of relaxation towards the fixed point given by *ν* _*D*_ = Λ_*I*_*/*2*π*. For excitatory coupling, the real eigenvalue can become positive with an associated saddle-node bifurcation and the emergence of collective chaos [64, 65].

By following [57], the Hopf boundaries can be identified by setting Λ = *i*2*πν* _*O*_ in (11) and to zero the real and imaginary part of the resulting equation, namely one gets

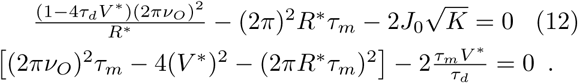

### B. Phase Diagrams of the Mean-Field Model

Apart from the linear stability of the asynchronous state and the associated Hopf boundaries which can be worked out analytically, the limit cycle solutions of the MF model and the associated bifurcations have been obtained by employing the software XPP AUTO developed for orbit continuation [66]. The MF model (8), apart from the membrane time constant *τ*_*m*_, which sets the system time scale, and the median in-degree *K*, which we fixed to 1000, is controlled by four independent parameters: namely, Δ_0_, *J*_0_, *I*_0_, *τ*_*d*_. In the following we will give an overview of the possible behaviours of the MF model in terms of two parameters phase diagrams for the most relevant combinations of the four mentioned parameters. The results of these analysis are summarized in Figs. 2 and 3.

**FIG. 2.**
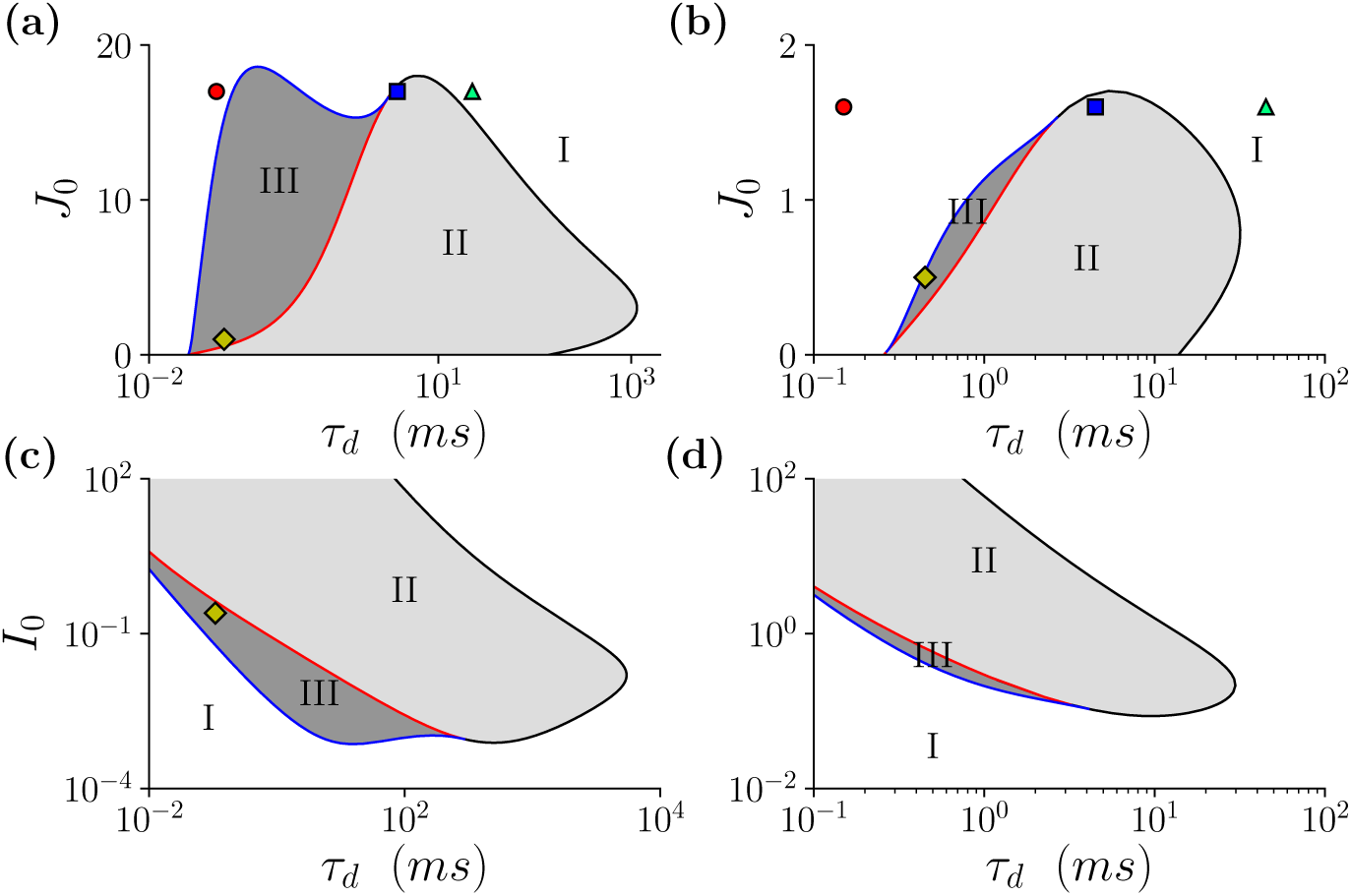
Phase diagrams of the mean-field model in the (*τ*_*d*_, *J*_0_)-plane (a-b) and in the (*τ*_*d*_, *I*_0_)-plane (c-d). The left panels refer to Δ_0_ = 0.3 and the right ones to Δ_0_ = 3. The red (black) line corresponds to sub-critical (super-critical) Hopf bifurcations, while the blue curve indicates saddle-node bifurcations of limit cycles. In the region I (white) the only stable solutions are foci and in the region II (light shaded) these are limit cycles. The dark shaded area (III) represents the region of coexistence of stable foci and limit cycles. The colored symbols indicate the states analyzed in Section IV. The parameters are *I*_0_ = 0.25 in (a-b) and *J*_0_ = 1.0 in (c-d) and *K* = 1000.

**FIG. 3.**
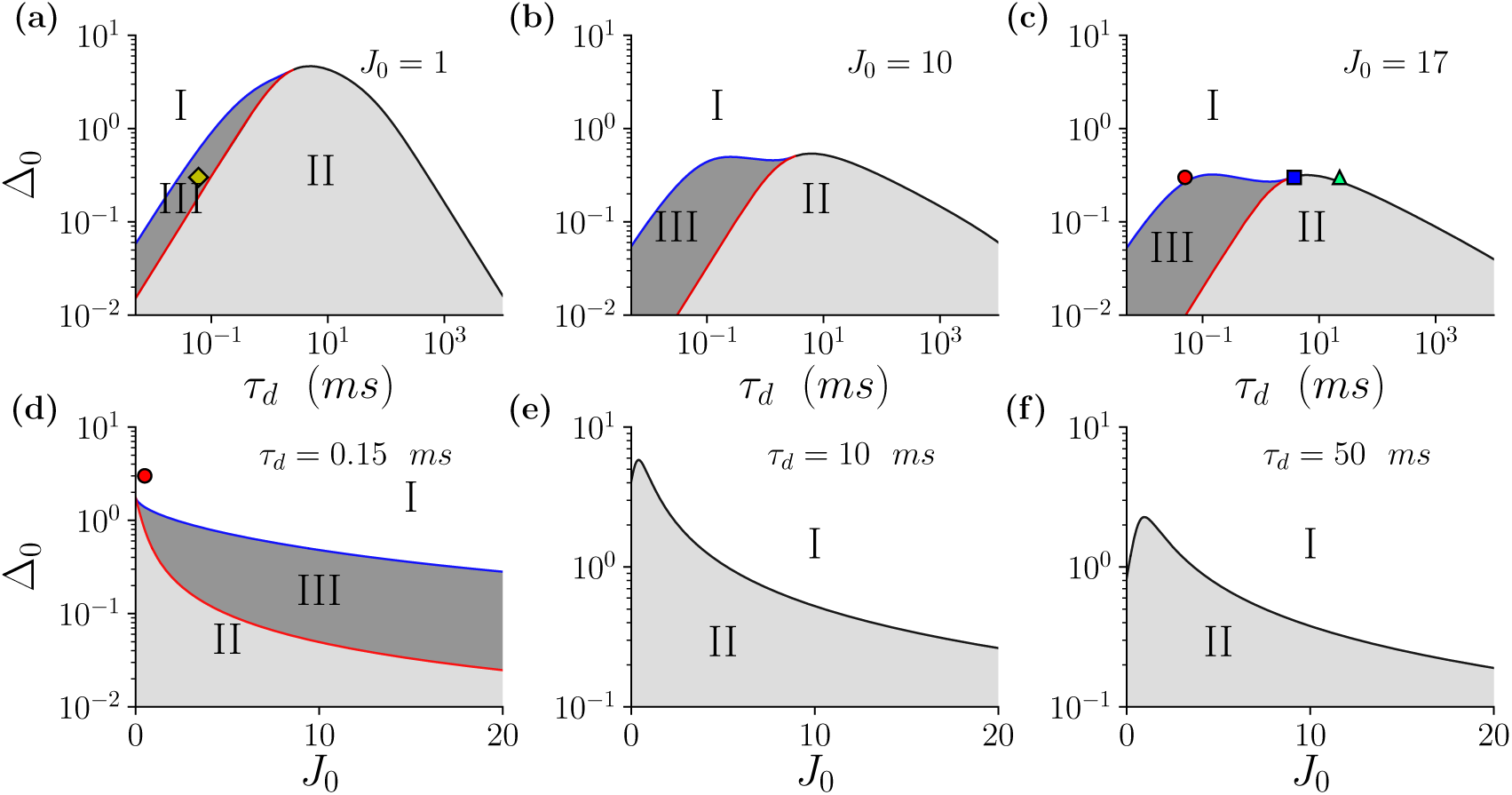
Phase diagrams of the mean-field model in the(*τ*_*d*_, Δ_0_)-plane (a-c) and in the (*J*_0_, Δ_0_)-plane (d-f) The line colors, the colored symbols and regions are defined as in Fig. 2. For the parameters we fixed *I*_0_ = 0.25 and K=1000.

Our analysis of the stationary solutions has revealed three possible regimes: stable foci (I); stable COs (II); co-existence of these two stable solutions (III). The stability boundaries of the COs are delimited by three kind of bifurcations: super-critical Hopf (black lines in the figures); sub-critical Hopf (red lines) and saddle-node (SN) of limit cycles (blue lines). Stable (unstable) COs emerge from stable foci at super-critical (sub-critical) Hopfs, while stable and unstable limit cycles merge at the SNs.

A fundamental parameter controlling the emergence of COs in the MF model is the synaptic time *τ*_*d*_, indeed in absence of this time scale no oscillations are present at the MF level [21]. On the other hand too large values of *τ*_*d*_ also lead to COs suppression, since the present model reduces to a Wilson-Cowan model for a single inhibitory population, that it is know to be unable to display oscillations [57]. As shown in Figs. 2 and 3, oscillations are observable for intermediate values of *τ*_*d*_ and not too large *J*_0_, since large inhibition leads to a quite reduced activity of the neurons not sufficient to ignite a collective behaviour. This is in agreement with the fundamental role played by gamma-Aminobutyric acid (GABA) in the emergence of epileptic seizures, characterized by an anomalous level of synchronization among the neurons, indeed the occurence of seizures seems strongly correlated with a GABA deficit, corresponding to a reduction of *J*_0_ in our case [67, 68]. Moreover, in order to observe COs the excitatory drive *I*_0_ should be larger than some critical value, as shown in Fig. 2 (c-d). This is consistent with the observation of the emergence of gamma oscillations in hippocampal slices induced through the acetylcoline agonist charbachol [50, 69], which leads to a decrease of the conductances of potassium channels, which can be mimicked as an increase of *I*_0_ [70, 71]. Indeed, by increasing the structural heterogeneity (measured by Δ_0_), which acts against coherent dynamics, larger values of *I*_0_ are required for COs as well as smaller synaptic couplings (see Figs. 2 (b),(d) and 3 (d-f)). Therefore the emergence of COs can be triggered by self-disinhibition as well as by an external excitatory drive, and we expect to observe in both cases the same scenarios.

As already mentioned, for infinitely fast synapses (*τ*_*d*_ → 0) the only possible solutions of the MF are foci characterized by two complex conjugate eigenvalues. Nevertheless, in the corresponding network the irregular firings of the neurons, due to the dynamical balance, can sustain COs, which are predicted to relaxed toward the fixed point in the MF. In the next Section we will analyze the role of these microscopic fluctuations in triggering the network dynamics also for finite *τ*_*d*_.

## IV. NETWORK DYNAMICS

We investigate in this Section the dynamics of the network by considering the parameter plan (*τ*_*d*_, *J*_0_). In particular, we want to examine the role of structural heterogeneity (measured by Δ_0_) in shaping the dynamical behaviours. This characteristic of the network structure is extremely relevant, as it can determine even if the system is in a balanced or in an imbalanced regime [21, 72, 73].

### A. High structural heterogeneity

We consider first a relatively high value for the structural heterogeneity, namely Δ_0_ = 3.0. For sufficiently large coupling *J*_0_, the bifurcation diagram reveals the emergence of oscillations in the MF model (8) via supercritical Hopf bifurcations, analogously to what has been reported for globally coupled networks [57]. An example of the bifurcation diagram, displaying the extrema of the mean membrane potential *V* as a function of *τ*_*d*_ is reported in Fig. 4 (a) for *J*_0_ = 1.6. In particular, we observe for instantaneous synapses (*τ*_*d*_ → 0) a stable focus, as expected from the analysis previously reported in [21]. The focus is stable up to 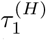 where it is substituted by a stable oscillatory state via a super-critical Hopf bifurcation. Oscillations are observable up to 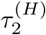, where via a second super-critical Hopf bifurcation they disappear and the unique stable solution for the MF system remains a focus. The typical stable regimes are denoted in Fig. 4 (a) by three capital letters: namely, (A) corresponds to a focus, (B) to a limit cycle and (C) to another focus. The network dynamics corresponding to these typical MF solutions is examined in the remaining panels of Fig. 4. For the focus solutions the network dynamics is asynchronous, as clearly visible from the corresponding raster plots in Fig. 4 (b) and (d). Furthermore, the dynamics of the neurons is quite regular in this case, as testified from the values of the average coefficients of variation, namely *CV* ≃ 0.14 and *CV* ≃ 0.04 corresponding to the distributions reported in Fig. 4 (e) and (f), respectively. At intermediate values of *τ*_*d*_, as predicted by the MF analysis, we observe COs with frequency *ν*_*OSC*_ ≃ 34 Hz in the network dynamics, see Fig. 4 (c). However, also in this case the dynamics is dominated by supra-threshold neurons with an associated very low *CV*, as evident from the large peak present at *cv*_*i*_ ≃ 0 in the distribution *P* (*cv*_*i*_) shown in Fig. 4 (g).

**FIG. 4.**
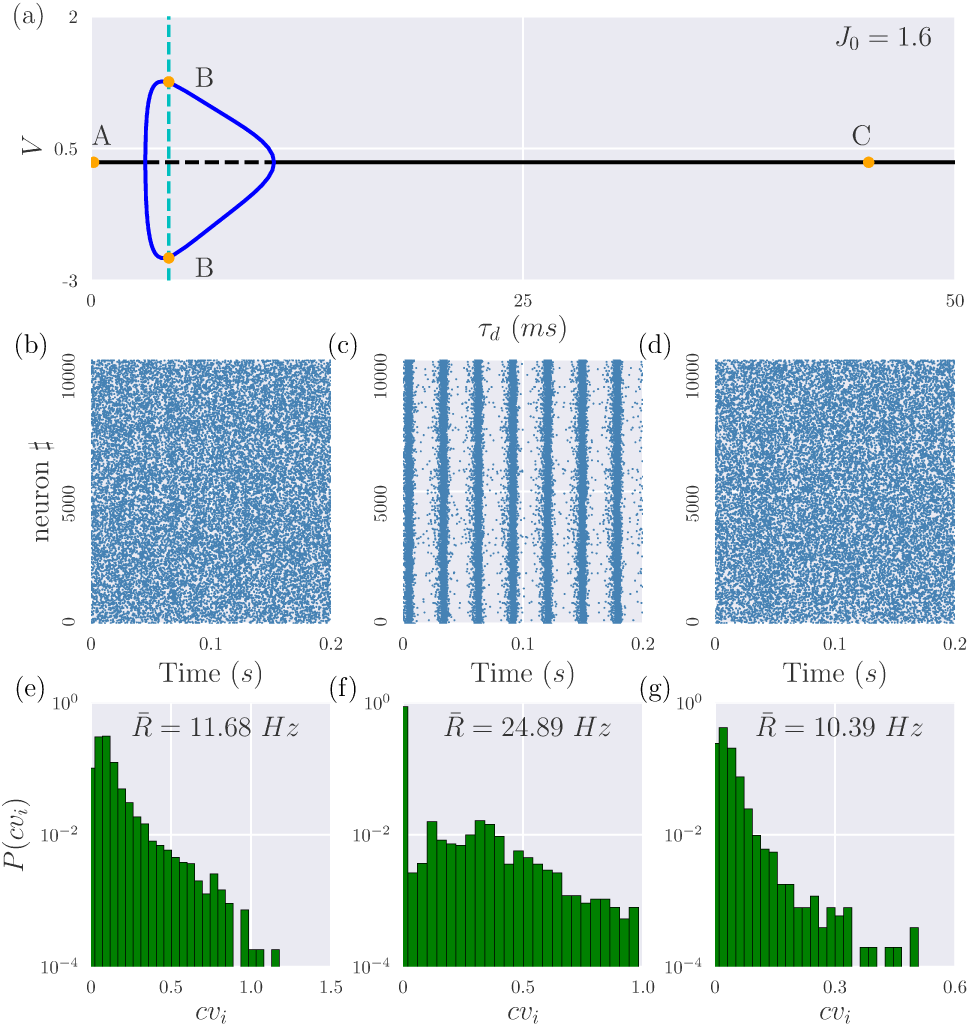
High Structural heterogeneity: super-critical Hopf bifurcation. (a) Bifurcation diagram of the MF model (8) displaying the extrema of *V* versus *τ*_*d*_, black solid (dashed) lines refer to the stable (unstable) focus, while blues solid lines to the oscillatory state. The supercritical Hopf bifurcations take place for 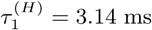 and 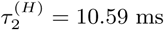. The capital letters in (a) denote three stationary states corresponding to different synaptic time scales, namely: (A) *τ*_*d*_ = 0.15 ms; (B) *τ*_*d*_ = 4.5 ms and (C) *τ*_*d*_ = 45 ms. The network dynamics corresponding to these states is reported in the panels below: the left column corresponds to (A), the central to (B) and the right one to (C). For each column, the top panels are the corresponding raster plots (b,c,d) and the bottom ones the distributions of the {cv_*i*_} of the single neurons (e,f,g). Network parameters are *N* = 10000, *K* = 1000 and Δ_0_ = 3.0. Other parameters are *I*_0_ = 0.25 and *J*_0_ = 1.6. The states (A), (B) and (C) are denoted in Fig. 2 (b) as a red circle, a blue square, and a green triangle, respectively.

For lower synaptic coupling *J*_0_ the phase portrait changes, as shown in Fig. 5 (a) for *J*_0_ = 0.5. In this case the MF analysis indicates that the transition from a stable focus to the oscillatory state occurs by increasing *τ*_*d*_ via a sub-critical Hopf bifurcation. At large synaptic coupling, the stable focus is recovered via a super-critical Hopf bifurcation taking place at 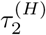, analogously to what has been seen for larger coupling. An interesting regime is observable between *τ* ^(*S*)^, where the stable and unstable limit cycle merge via a saddle-node bifurcation, and 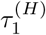, where the focus become unstable. In this interval the MF model displays two coexisting stable solutions: a limit cycle and a focus. It is important to verify if also the finite size sparse network displays this coexistence, indeed as shown in Fig. 5 depending on the initial conditions the network dynamics can converge towards COs or towards an asynchronous state. In particular, we observe that the asynchronous dynamics is associated to extremely low *cv*-values (see Fig. 5 (d)) suggesting that this can be considered as a sort of irregular splay state [74]. However, also the COs with *ν*_*OSC*_ ≃ 58 Hz are characterized by a low average coefficient of variation, namely *CV* ≃ 0.014 indicating that the dynamics is mean driven. The sub-critical Hopf, as expected, is associated to a hysteretic behaviour, this effect can be revealed by considering simulations concerning an adiabatic variation of *τ*_*d*_. The results of these simulations are reported in Fig. 5 (b), where the maximal values of the instantaneous firing rate *R*_*M*_ are reported as a function of *τ*_*d*_ for the adiabatic protocol and compared with the MF estimations of *R*_*M*_. From the figure it is clear that the transition from the focus to the stable limit cycle occurs at 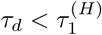 and the system returns from the oscillatory state to the asynchronous one at *τ*_*d*_ definitely smaller than *τ* ^(*S*)^. These are finite size (and possibly also finite time) effects, indeed as shown in Fig. 5 (b) by increasing *N* the transition points approach the MF ones.

**FIG. 5.**
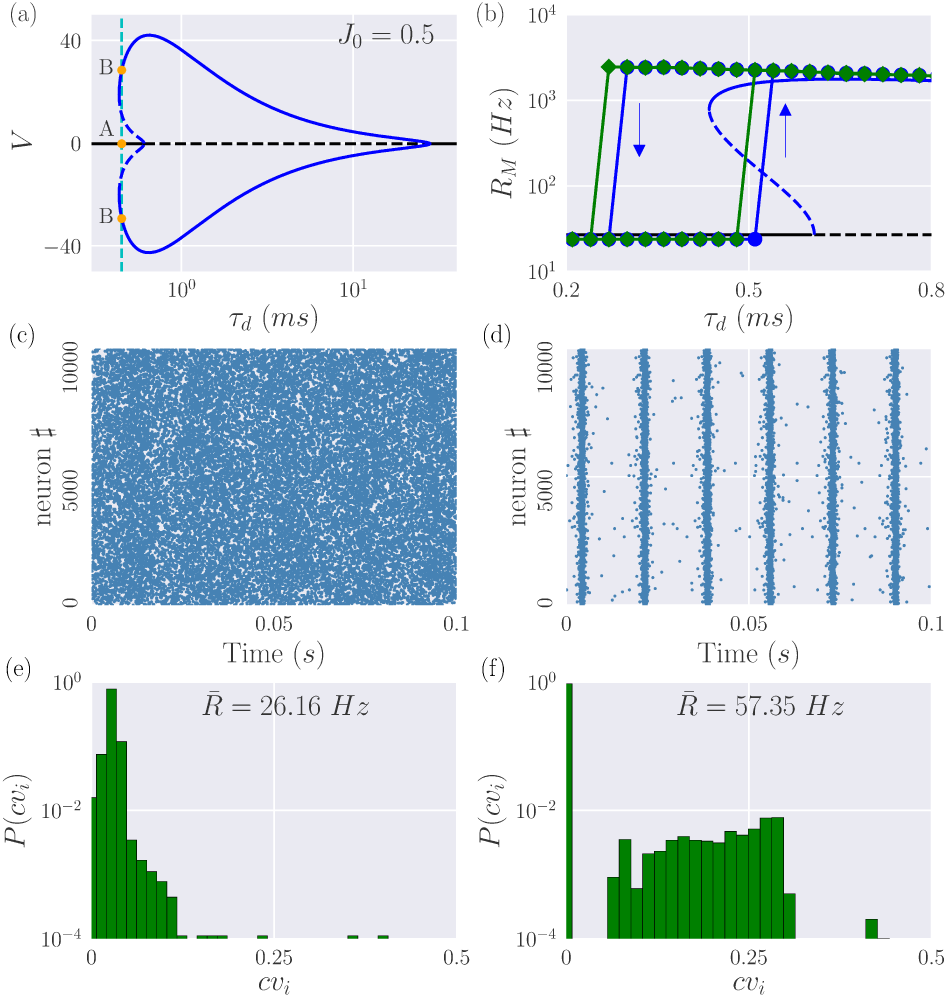
High structural heterogeneity: sub-critical Hopf bifurcation. (a) Bifurcation diagram of the MF model analogous to the one reported in Fig. 4 (a). The super-critical (sub-critical) Hopf bifurcation takes place at 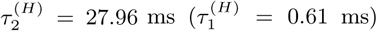, the saddle node of limit cycles at *τ* ^(*S*)^ = 0.43 ms. The capital letters in (a) denote two stationary states corresponding to the same synaptic time scales *τ*_*d*_ = 0.45 ms. This state is denoted in Fig. 2 (b) as a yellow diamond. The network dynamics corresponding to these states is reported in the panels below: the left column correspond to (A) and the right one to (B). For each column, the top panels display the raster plots (c,d) and the bottom ones the distribution of the *{cv*_*i*_*}* of the single neurons (e,f). In panel (b) are reported the maximal values of the rate *R*_*M*_ obtained by performing adiabatic simulations by first increasing and then decreasing the synaptic time *τ*_*d*_ (green) diamonds for *N* = 10, 000 and (blue) circles for *N* = 20, 000, the arrows denote the jump from one state to the other. The MF results are also displayed: solid (dashed) black lines refer to stable (unstable) foci, while solid (dashed) blue lines to stable (unstable) limit cycles. Parameters are the same as in Fig. 4, apart for *J*_0_ = 0.5, the parameters for the adiabatic simulations are Δ*τ*_*d*_ = 0.03 ms, 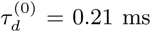 and 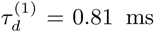.

### B. Low structural heterogeneity

We consider now a relatively low value of the structural heterogeneity, i.e. Δ_0_ = 0.3, which for instantaneous synapses can sustain a dynamically balanced state [21]. Let us first consider a relatively large coupling, namely *J*_0_ = 17.0, the corresponding bifurcation diagram for the MF model is reported in Fig. 6(a). This is quite similar to the one previously shown for high structural heterogeneity in Fig. 4(a). However, peculiar differences are observable at the level of network simulations. Indeed in this case COs are present for all the considered *τ*_*d*_-values, even if these correspond to stable foci in the MF (states (A) and (C) in Fig. 6(a)) as evident from the raster plots reported in Fig. 6(b) and (d). In particular we measured the following frequencies for the observed COs: *ν*_*OSC*_ ≃ 57 Hz for state (A), *ν*_*OSC*_ ≃ 30 Hz for (B) and *ν*_*OSC*_ ≃ 16 Hz for (C). Furthermore, the network dynamics is now definitely more irregular than for high Δ_0_ with distributions *P* (*cv*_*i*_) centered around *cv*_*i*_ = 1 for the states (A) and (C) in Fig. 6(a) corresponding to stable foci in the MF formulation (see Fig. 6(e) and (g)) and with *P* (*cv*_*i*_) extending towards values around *cv*_*i*_ ≃ 1 for the oscillatory state (B), as shown in Fig. 6(i). This irregularity in the spike emissions is a clear indication that now the dynamics is mostly fluctuation driven due to the dynamically balanced dynamics observable in the sparse network for sufficiently low structural heterogeneity. Furthermore, as shown in [21] for instantaneous synapses, these current fluctuations are able to turn the macroscopic damped oscillations towards the stable foci, observable in the MF model, in sustained COs in the network. The origin of the COs observable for the state (B) is indeed different, since in this case sustained oscillations emerge due to a super-critical Hopf bifurcation both in the MF and in the network dynamics.

**FIG. 6.**
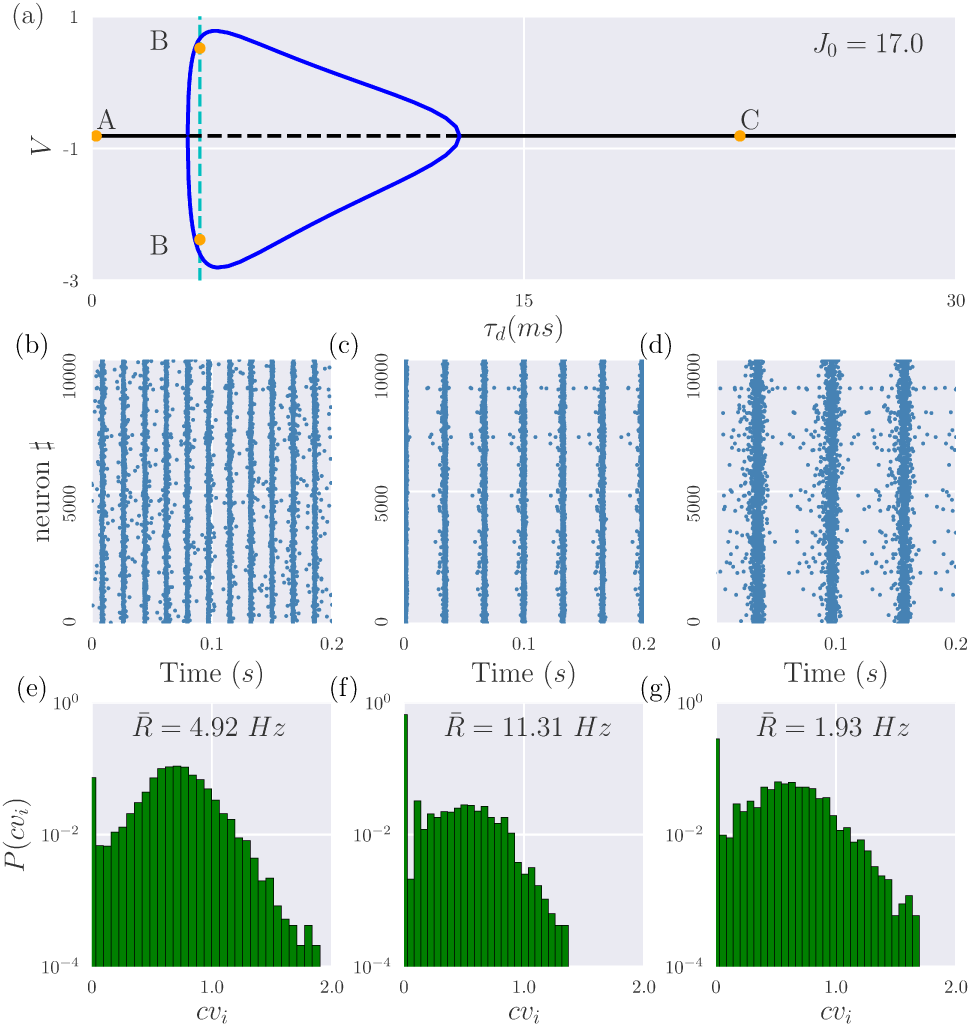
Low structural heterogeneity: super-critical Hopf bifurcation. The panels here displayed are analogous to the ones in Fig. 4. In this case the super-critical Hopf bifurcations occur for 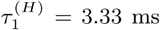 and 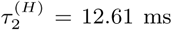 and the stationary states in (a) corresponding to the capital letter (A), (B) and (C) refer to *τ*_*d*_ = 0.15 ms, *τ*_*d*_ = 3.75 ms and *τ*_*d*_ = 22.5 ms, respectively. The parameters are the same as in Fig. 4, apart Δ_0_ = 0.3. and *J*_0_ = 17. The states (A), (B) and (C) are denoted in Fig. 2 (a) and Fig. 3 (c) as a red circle, a blue square, and a green triangle, respectively.

By decreasing the synaptic coupling *J*_0_ (Fig. 7(a)) we observe in the MF phase diagram the emergence of regions where the oscillations coexist with the stable focus in proximity of a sub-critical Hopf bifurcation, analogously to what has been reported for high heterogeneity (see Fig 5(a)). At variance with that case, we have now in the network a bistability between two COs whose origin is different: one emerges via a Hopf bifurcation and it is displayed in Fig. 7(d), while the other is sustained by the irregular spiking associated to the balanced state and the corresponding raster plot is reported in Fig. 7(c). In particular, the latter COs are associated to large *cv*-values (Fig 7 (e)) typical of a balanced regime, while the other COs are extremely regular as shown in Fig 7 (f) resembling the dynamics of a highly synchronized system.

**FIG. 7.**
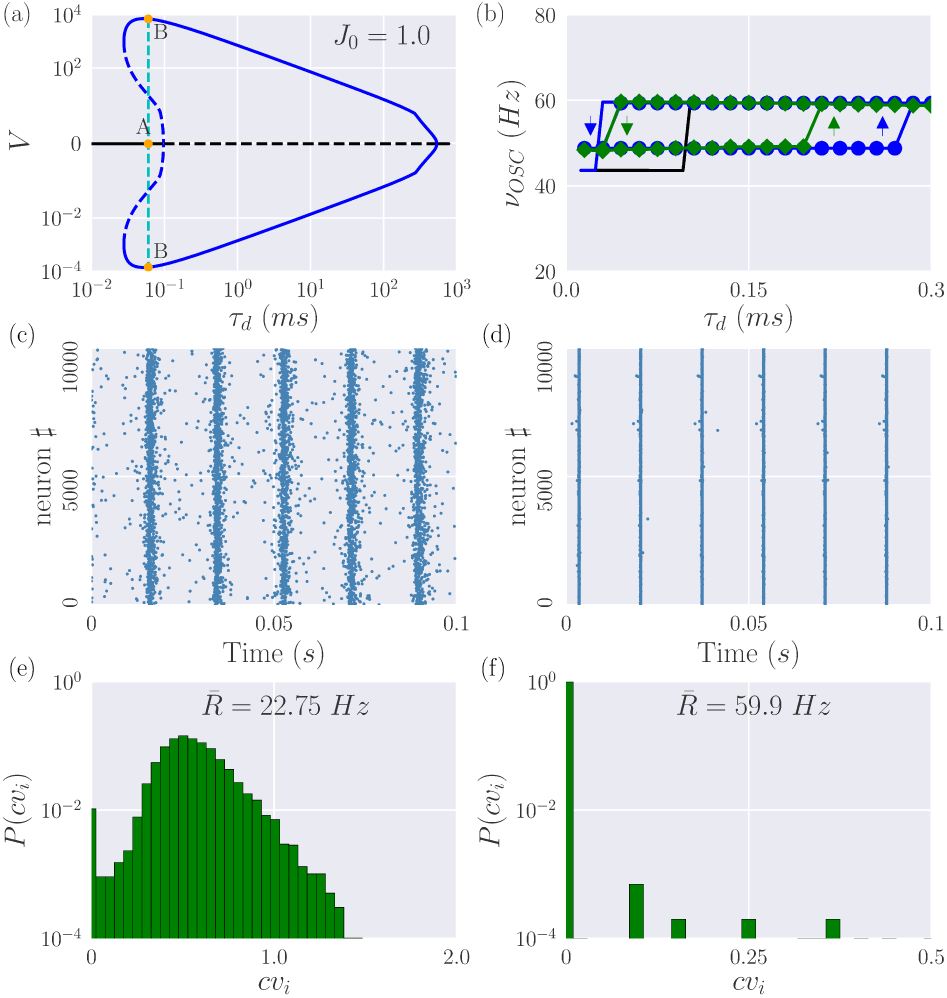
Low structural heterogeneity: sub-critical Hopf bifurcation. The panels here displayed, apart panel (b), are analogous to the ones in Fig. 5. In the present case the sub-critical Hopf occurs at 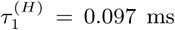, while the super-critical Hopf at 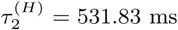 and the saddle-node of limit cycles at *τ* ^(*S*)^ = 0.028 ms, and the coexisting states (A) and (B) shown in (a) refer to *τ*_*d*_ = 0.06 ms. This state is denoted in Figs. 2 (a) and (c) and Fig. 3 (a) as a yellow diamond. Panel (b) reports the frequency of collective oscillations as measured via adiabatic simulations for *N* = 2000 by considering *T*_*s*_ = 90 ms (blue circles) and *T*_*s*_ = 1500 ms (green diamonds), the transient time *T*_*t*_ = 15 ms is unchanged. The solid lines in (b) refer to the MF results, namely the black line to *ν*_*D*_ and the blue one to the limit cycle frequency *ν*_*O*_. The parameters are as in Fig. 6 apart for *J* = 1.0 and for the adiabatic simulations are Δ*τ*_*d*_ = 0.015 ms, 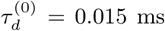 and 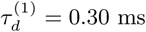.

In order to analyze the coexistence region, we report in Fig. 7(b) the frequencies *ν*_*OSC*_ of the collective oscillations as measured via adiabatic simulations of the network (symbols). Furthermore, the MF results for *ν*_*D*_ associated to the foci and the frequencies *ν*_*O*_ of the limit cycles are also reported in the figure as black and blue solid line, respectively. The frequencies of the COs in both states are reasonably well captured by the MF approach, furthermore the two frequencies can be quantitatively associated to fast and slow gamma oscillations. The comparison reveals that the COs induced by microscopic irregular firing exist far beyond 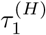, despite here the unique stable solution predicted by the MF should be the almost synchronized bursting state shown in Fig. 7(d). On the other hand the backward transition is almost coincident with the MF prediction for *τ* ^(*S*)^ as displayed in Fig. 7(b). As reported in Fig. 7(b), we observe that also the forward transition value approaches to 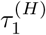 by increasing the duration *T*_*s*_ of the adiabatic steps. Therefore this result suggest that the observed discrepancies are due to finite time (and possibly finite size) effects affecting the network simulations.

## V. COEXISTENCE OF SLOW AND FAST GAMMA OSCILLATIONS

In the previous Section we have shown, for a specific choice of the parameters, that fast and slow collective gamma oscillations can coexist. However, the phenomenon is observable in the whole range of coexistence of the stable foci and of the stable limit cycles. In particular, in Fig. 8 we report in the (*τ*_*d*_, *J*_0_)-plane the frequencies *ν*_*D*_ associated to the damped oscillations towards the MF focus in panel (a) and the frequencies *ν*_*O*_ of the limit cycles in panel (b). It is evident that *ν*_*D*_ ≃ 30 − 40 Hz, while the frequencies of the limit cycle *ν*_*O*_ are of the order of 60 Hz, thus in the network we expect to observe coexisting COs characterized by slow and fast rhythms in a wide range of parameters.

**FIG. 8.**
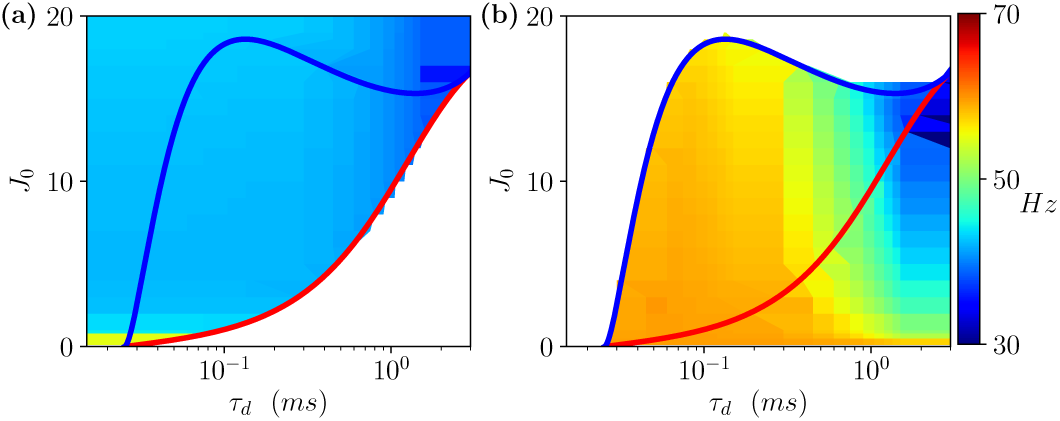
Coexisting fast and slow gamma oscillations. (a) Frequencies *ν*_*D*_ associated to the damped oscillations towards the stable foci; (b) frequencies *ν*_*O*_ of the limit cycles. Red lines refer to the sub-critical Hopf boundaries, while the blue ones to saddle-node bifurcations of limit cycles. Other parameters are *I*_0_ = 0.25 and Δ_0_ = 0.3.

For this parameter set *ν*_*D*_ seems to depend only slightly on *τ*_*d*_ and *J*_0_. On the contrary the frequency *ν*_*O*_, characterizing the more synchronized events, is influenced by these parameters. In particular, *ν*_*O*_ decreases for increasing IPSP time duration, similarly to what observed experimentally for cholinergic induced gamma oscillations in the hippocampus *in vitro* [50]. Moreover, barbiturate, a drug often used as anxiolytic, is known to increase IPSP time duration [75] and slow down gamma oscillations [76], in accordance with our scenario. Furthermore, for *τ*_*d*_ > 1 ms the increase of *J*_0_ leads to a decrease of *ν*_*D*_, similarly to the effect of alcohol that induces an increase of inhibition associated to a decrease in gamma oscillation frequencies measured in the human visual cortex [77].

The coexistence of fast and slow gamma COs is a quite general phenomenon not limited to the specific network topology we employed, i.e. that associate to the Lorentzian in-degree distribution. Indeed, as shown in Appendix A it can be observed also for a sparse Erdös-Renyi network.

### A. Switching gamma rhythms

As a further aspect, we will consider the possibility to develop a simple protocol to drive the system from slow gamma COs to fast ones (and vice versa) in the bistable regime. Let us consider the case where the network is oscillating with slow gamma frequency as shown in Fig. 9 for *I*_0_ ≡ *I*_1_ = 0.25. The protocol to drive the system in the fast gamma band consists in delivering a step current *I*_2_ to all the neurons for a very limited time interval *T*_*sh*_. In this way the system is transiently driven in a regime where oscillatory dynamics is the only stable solution, as a matter of fact the neurons remain in a high frequency state even after the removal of the stimulation, when *I*_0_ returns to the initial value *I*_1_ (see Fig. 9). In order to desynchronize the neurons and to recover the slow gamma COs, we delivered random quenched DC currents *I*_0_(*i*) (with *i* = 1, …, *N*) to the neurons for a time period *T*_*sl*_. The currents *I*_0_(*i*) are taken from a flat distribution with a very low average value *I*_3_ and a width Δ*I*_3_ corresponding to a parameter range where the MF foci are the only stable solutions. As shown in Fig. 9 in this case to drive the system from fast to slow gamma oscillations it was sufficient to apply the perturbation for a much smaller period *T*_*sl*_ ≪ *T*_*sh*_.

**FIG. 9.**
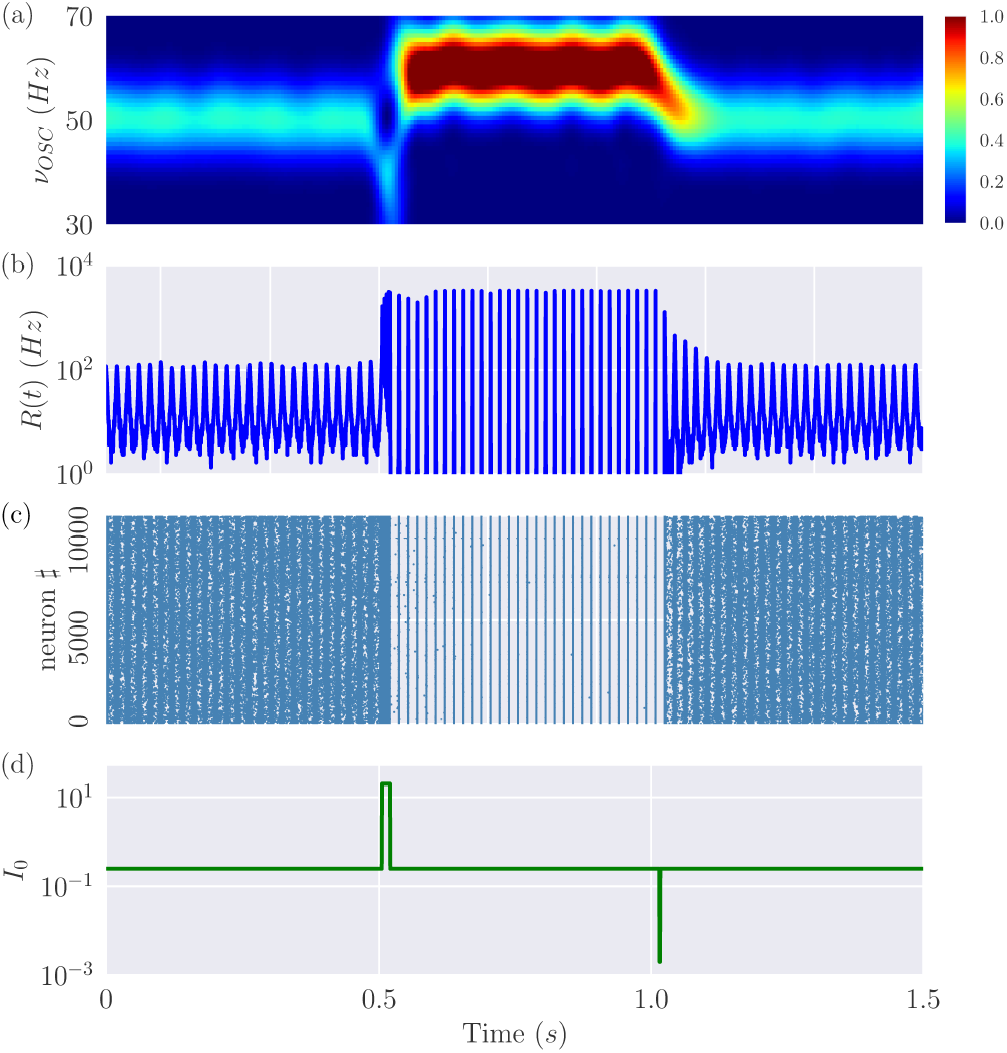
Switching from fast (slow) to slow (fast) gamma oscillations. Results of the switching experiments described in the text, from top to bottom: (a) spectrogram of the mean membrane potential *V* ; (b) the firing rate *R*(*t*); (c) the raster plot and (d) the stimulation protocol reporting the average external DC current. The parameters are the same as in Fig. 7 (in particular *τ*_*d*_ = 0.06 ms), apart *T*_*sh*_ = 0.015 s, *T*_*sl*_ = 0.0015 s, *I*_1_ = 0.25, *I*_2_ = 20.0, *I*_3_ = 0.012, Δ*I*_3_ = 0.01.

Let us now try to characterize in more details the observed switching transitions. This can be done by considering the MF bifurcation diagram in terms of the external DC current *I*_0_ reported in Fig. 10 (a) for the examined parameters. The diagram reveals a sub-critical Hopf bifurcation taking place at *I*^(*H*)^ ≃ 0.43 and a region of bistability extending from *I*^(*S*)^ ≃ 0.06 to *I*^(*H*)^. Therefore, if we consider a DC current in the bistable interval (namely, *I*_0_ ≡ *I*_1_ = 0.25) and we prepare the system in the slow gamma regime a transition to the fast gamma COs will be observable whenever the DC current is increased to a value *I*_0_ ≡ *I*_2_ > *I*^(*H*)^. However, if we return in the bistable regime at *I*_0_ ≡ *I*_1_, after delivering the perturbation *I*_2_ for a time interval *T*_*P*_, it is not evident in which regime (fast or slow) the system will end up. Thus we have measured the transition probability from slow to fast gamma for different *T*_*P*_ and *I*_2_ by following the protocol reported in Section II B. We analized these transitions in presence of a small additive noise on the membrane potentials of amplitude *A*_*n*_, somehow en-compassing the many sources of noise present in neural circuits.

**FIG. 10.**
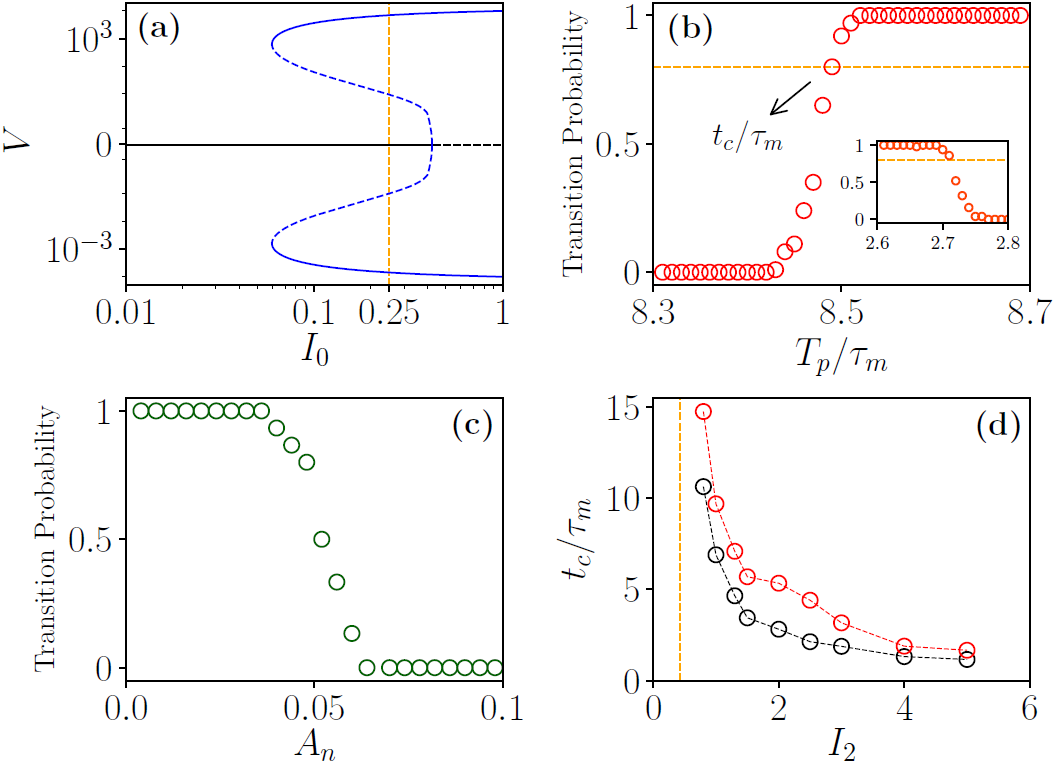
Statistics of the switching transitions. (a) Bifurcation diagram of the MF model reporting the extrema of the mean membrane potential *V* as a function of *I*_0_ displaying stable (solid line) and unstable solutions (dashed lines) for foci (black) and limit cycles (blue). The vertical dashed (orange) line refers to *I*_0_ = 0.25. (b) Transition probability as a function of *T*_*P*_, the orange dashed line denotes the 80 % for *I*_2_ = 1.0, in the inset the data for the transition from fast to slow gamma is reported for *I*_3_ = 0.03, in both cases *A*_*n*_ = 0.05 (c). Transition probability as a function of the noise amplitude *A*_*n*_ for *I*_2_ = 1 and *T*_*P*_ = 8.48*τ*_*m*_. (d) Crossing times *t*_*c*_ versus the perturbation amplitude *I*_2_ for various noise levels: *A*_*n*_ = 0.02 (black circles) and 0.07 (red circles). The vertical orange line indicates the value *I*^(*H*)^. Panels (c-d) refer to the transition from slow to fast gamma COs, while the inset in (b) to the transition from fast to slow gamma. The parameters are the same as in Fig. 7.

The results shown in Fig. 10 (b) for *I*_2_ = 1.0 and *A*_*n*_ = 0.05 reveal that even if *I*_2_ > *I*^(*H*)^ the perturbation should be applied for a minimal time interval *T*_*P*_ > *t*_*c*_ ≃ 0.12 s to induce the transition to the fast gamma COs in at least the 80% of cases. It is interesting to note that the noise amplitude can play a critical role on the switching transition, indeed the increase of *A*_*n*_ can desynchronize the fast gamma regime even for *T*_*P*_ > *t*_*c*_, see Fig. 10 (c). Therefore *t*_*c*_ depends critically not only on *I*_2_ but also on *A*_*n*_: as expected by increasing *I*_2_ the crossing time drops rapidly towards zero, while the switching transition is delayed to longer times for larger *A*_*n*_ (see Fig. 10 (d)).

For what concerns the transition from fast to slow gamma, this occurs in an irreversible manner only for amplitude of the perturbation *I*_3_ < *I*^(*S*)^, an example is shown in the inset of Fig. 10 (b). Despite the switching transition can be observed also for *I*_3_ > *I*^(*S*)^ this will be much more complex due to the competition of the two stable states in the interval *I*_0_ ∈ [*I*^(*S*)^ : *I*^(*H*)^] and more specific protocols should be designed to obtain the desynchronization of the system.

### B. Theta-gamma cross-frequency coupling

So far we have described a simple protocol where external constant stimulations to the inhibitory network can drive the neural population from one state to the other. However, gamma oscillations are usually modulated by theta oscillations in the hippocampus and in the neocortex during movement and REM sleep [34, 44]. This has recently inspired a series of optogenetic experiments *in vitro* intended to reproduce the effect of the theta forcing and the activity observed *in vivo* [39, 48, 49]. To make a closer contact with these experiments we decided to consider a periodic stimulation of all neurons in the network as follows:

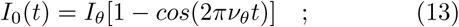

where the phase of the theta forcing is defined as *θ*(*t*) = 2*πν*_*θ*_*t*. The term appearing in (13) corresponds to the synaptic input received by the neurons, in order to compare this forcing term with the LFPs experimentally measured in [35, 46] and which reveals theta oscillations, one should remember that the LFP corresponds to the electrical potential measured in the extracellular medium around neurons [78]. Therefore for a meaningful comparison with the synaptic input (13) the sign of the LFP should be reversed. This is consistent with the observations reported in [35, 46] that the maximum of activity of the excitatory (pyramidal) cells is observed in correspondence of the minimum of the LFP.

We considered the network dynamics in presence of the periodic forcing (13) and additive noise on the membrane potentials (with zero mean and amplitude *A*_*n*_). As shown in Fig. 11, the response of the system to the forcing is controlled by the value of the amplitude *I*_*θ*_ in (13): for small *I*_*θ*_ ≤ 0.20 one observes only slow gamma COs; for intermediate values of the amplitude 0.20 < *I*_*θ*_ ≤ 0.32 one has the coexistence of slow and fast gamma COs; while for *I*_*θ*_ ≥ 0.32 only fast oscillations are present.

**FIG. 11.**
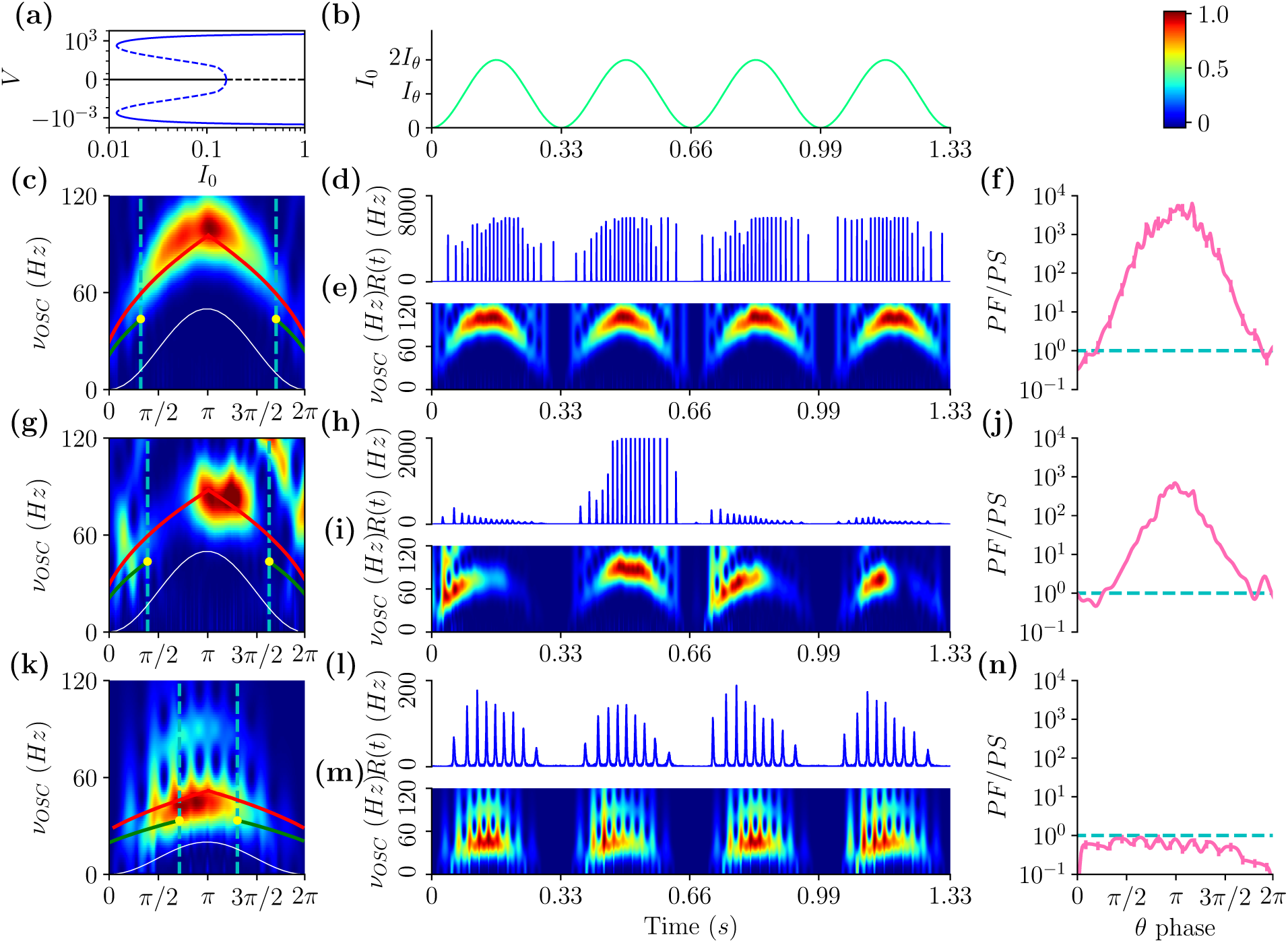
Fast and slow gamma oscillations entrainment with the theta forcing. (a) Bifurcation diagram of the MF model analogous to the one reported in Fig. 10 (a). (b) Theta forcing (13) versus time. The three lower rows refer from top to bottom to *I*_*θ*_ = 0.35, 0.3 and 0.1 with noise amplitudes *A*_*n*_ = 1 for the two largest values and *A*_*n*_ = 1 × 10^−3^ for the smallest one. In the left column are reported the averaged spectrograms as a function of the theta phase. In the same panels are reported *ν*_*D*_ (solid green line), *ν*_*O*_ (solid red line) as a function of *θ*, as well as the forcing in arbitrary units (white solid line). The vertical blue dashed lines indicate the stability boundaries for the focus solution: the focus is unstable for *θ* phases within such lines. The central column displays an instance over four theta cycles of the corresponding spectrograms and of the instantaneous firing rates *R*(*t*). The right column reports the ratio *PF/P S* of the power contained in the fast (50 *< ν*_*OSC*_ *<* 100 Hz) and slow (30 *≤ ν*_*OSC*_ ≤ 50 Hz) gamma bands as a function of the *θ* phase. In this case the error bars are displayed, but are almost invisible on the reported scale. Parameters are *J*_0_ = 1, *τ*_*d*_ = 0.15 ms, Δ = 0.3 and *K* = 1000, for the simulations we considered *N* = 10000, and *ν*_*θ*_ = 3 Hz, the data for the spectrograms (left row) have been obtained by averaging over 30 theta cycles and those for *PF/PS* (right row) over 400 cycles.

For small *I*_*θ*_, as one can appreciate from the raster plot in panel (m), the firings of the neurons are quite irregular and the bursts involve only a limited number of neurons, as confirmed also by the low values of *ρ* during the firing activity (see Fig. 12 (c)). Furthermore the corresponding spectrogram in Fig. 11 (k) reveals that the power is concentrated at frequencies below 50 Hz and that the amplitude of the spectrum has a modulated structure as a function of the phase. This is confirmed by the analysis of the power of the spectrum *PS* (*PL*) restricted to the slow (fast) gamma band (see Fig. 11 (n)). These are indications of theta-nested gamma oscillations, as confirmed by the instantaneous firing rate reported in Fig. 11 (l), which reveals also an evident P-A coupling between the gamma phases and the theta forcing. In this case we considered a quite small value of the amplitude *A*_*n*_ of the extrinsic noise, this in order to compare our findings with results obtained in *in vitro* experiments.

**FIG. 12.**
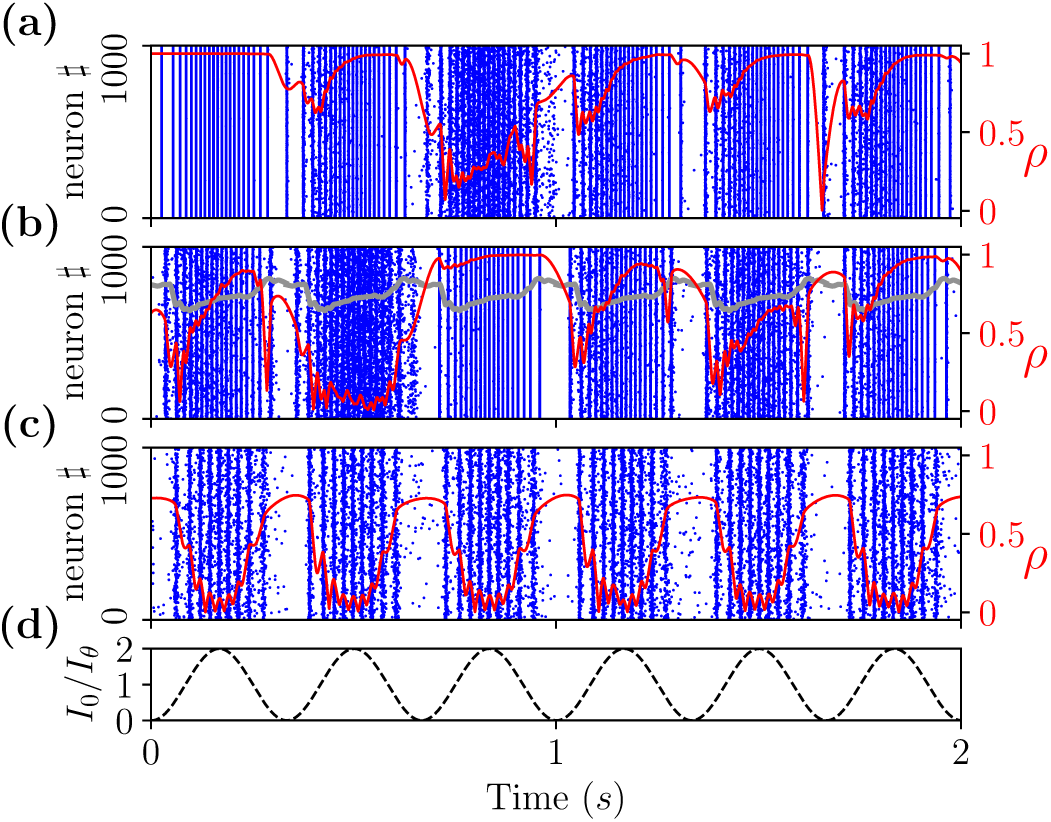
Level of synchronization of fast and slow gamma oscillations. (a-c) Raster plots and Kuramoto order parameter *ρ* (solid red curve) in presence of theta forcing (13), the data refer from top to bottom to *I*_*θ*_ = 0.35, 0.3 and 0.1 with amplitude of noise as in Fig. 11. The grey curve in panel (b) represents the value of *ρ* for one *θ*-period averaged over 50 *θ*-cycles. (d) Theta forcing profile versus time. Parameters are *J*_0_ = 1, *τ*_*d*_ = 0.15 ms, Δ = 0.3, *K* = 1000, *N* = 10000 and *ν*_*θ*_ = 3 Hz.

Indeed, these results resemble experimental observations of theta-nested gamma oscillations induced *in vitro* by sinusoidal optical stimulation at theta frequency in the medial entorhinal cortex (mEC) [49] and in the areas CA1 [39] and CA3 [48] of the hippocampus. In all these experiments single neurons spiked quite irregularly, while the collective dynamics was oscillatory, analogously to our dynamics as shown in Fig. 11 (l) and (m). As previously discussed, these COs are induced by intrinsic fluctuations and characterized at a MF level by frequencies ≃ *ν*_*D*_ (green solid line), which represents a reasonable estimation of the position of the maxima of the spectrogram as shown in Fig. 11 (k).

The situation is quite different for sufficiently large forcing amplitude, where the neuronal dynamics becomes quite regular and highly synchronized, as evident from Figs. 11 (d) and (e) and by the value of *ρ* reported in Fig. 12 (a). In this case the power is concentrated in the fast gamma band and it is maximal in correspondence of the largest value of *I*_0_ occuring at *θ* = *π* (see Figs. 11 (c) and (f)). Furthermore the profile of the maximal power in the spectrogram follows reasonably well the MF values *ν*_*O*_ (red solid line) expected for fast gamma COs, as evident from Fig. 11 (c). For these large currents we have a sort of pathological synchronization usually observable in connection with neuronal diseases. In particular, highly synchronized fast gamma oscillations have been observed in patients with neocortical epilepsy [79].

The most interesting situation occurs for intermediate amplitudes, specifically we considered *I*_*θ*_ = 0.30. As evident from Figs. 11 (h) and (i) in this case the network dynamics can vary noticeably from one theta cycle to the next, due to the switching from one gamma regime to the other occurring erratically induced by the larger extrinsic noise here employed (namely, *A*_*n*_ = 1) to mimic *in vivo* conditions. This is particularly evident in Fig. 12 (b), where the network dynamics reveals a large variability as testified by the level of synchronization from one cycle to another. However by averaging over a sufficiently large number of cycles we can identify stationary features of this dynamics. In particular, we observe that the values of maximum power in the averaged spectrum correspond to different theta phases for the slow and fast gamma COs: namely, for slow gamma the maximal activity is observable at small angles, while for fast gamma this corresponds to the largest value of the forcing current (13) (see Figs. 11 (g) and (j)). Furthermore, on average we observe a steady increase of *ρ* within a *θ* cycle (see the grey line in Fig. 12 (b)), confirming that slow irregular gamma COs are mostly observable at small angles, while fast more regular gamma COs set in at *θ ≥ π*.

These findings resemble the experimental results reported in [46] for the region CA1 of the hippocampus in freely moving rats, where it has been reported that slow gamma power were peaked around *θ* ≃ 0.4*π* and fast gamma power around *θ* ≃ *π*, corresponding also to the maximum of activity of excitatory place cells. Similar results have been reported in [35] for what concerns the slow gamma rhythm, however in those experiments fast gamma (referred in as intermediate gamma) occurs earlier in the theta cycle. Recent experiments obtained on behaving rats singled out a wide cycle-to-cycle variability of the frequency of CA1 theta-nested gamma oscillations and an overlap of the frequency bands associated to slow and fast gamma oscillations [80]. These results do not contradict previous findings which were mainly based on spectral features obtained by averaging over many cycles [35, 46]. Our results confirm the absence of this contradiction. Indeed, the instantaneous spectrograms reported in Fig. 11 (i) show a large cycle-to-cycle variability, while the average spectrogram shown in Fig. 11 (g) displays distinct peaks for slow and fast gamma COs at specific *θ*-angles.

The network response to the external periodic forcing (13) can be interpreted in terms of an adiabatic variation of the external current whenever the time scale of the forcing term is definitely slower with respect to the neuronal time scales (i.e. *τ*_*m*_ and *τ*_*d*_). Since this is the case, we can try to understand the observed dynamics at a first level of approximation by employing the bifurcation diagram of the MF model obtained for a constant DC current *I*_0_, which is shown in Fig. 11 (a) for the set of parameters here considered. The diagram reveals that the system bifurcates via a sub-critical Hopf from the asynchronous state to regular oscillatory behaviour at a current *I*^(*H*)^ ≃ 0.159 and that the region of coexistence of stable foci and limit cycles is delimited by a saddle-node bifurcation occurring at *I*^(*S*)^ ≃ 0.012 and by *I*(*H*).

The forcing current (13) varies over a theta cycle from a value *I*_0_ = 0 at *θ* = 0 up to a maximal value *I*_0_ = 2*I*_*θ*_ at *θ* = *π* and returns to zero at *θ* = 2*π*. Since the forcing current will start from a zero value, we expect that the network will start oscillating with slow gamma frequencies associated to the stable focus which is the only stable solution at small *I*_0_ < *I*^(*S*)^. Furthermore, if *I*_*θ*_ < *I*^(*H*)^*/*2 the system will remain always in the slow gamma regimes during the whole forcing period, since the focus is stable up to the current *I*^(*H*)^.

For amplitudes *I*_*θ*_ > *I*^(*H*)^/2 we expect a transition from slow to fast COs for a theta phase *θ*^(*H*)^ = arccos [(*I*_*θ*_ − *I*^(*H*)^)/*I*_*θ*_] corresponding to the crossing of the sub-critical Hopf. Since this transition is hysteretic the system will remain in the fast regime until the forcing current does not become extremely small, namely *I*_0_ *< I*^(*S*)^, corresponding to a theta phase *θ* ^(*S*)^ = 2*π* − arccos [(*I*_*θ*_ − *I*^(*S*)^)*/I*_*θ*_].

The performed analysis is quasi-static and does not take into account the time spent in each regime. If *I*_*θ*_ >> *I*^(*H*)^ the time spent by the system in the slow gamma regime is extremely reduced, because *θ*^(*H*)^ ≃ 0 and *θ*^(*S*)^ ≃ 2*π*, and this explains why for large *I*_*θ*_ we essentially observe only fast gamma. On the other hand, we find only slow gamma COs for *I*_*θ*_ up to 0.20, a value definitely larger than *I*^(*H*)^/2, and this due to the fact that a finite crossing time is needed to jump from one state to the other as discussed in the previous sub-section.

Let us now focus on the case *I*_*θ*_ = 0.3, where we observe the coexistence of fast and slow gamma COs. As already mentioned we have stable foci in the range *I*_0_ ∈ [0 : *I*^(*H*)^], this in terms of *θ*-angles obtained via the relationship (13) for *I*_*θ*_ = 0.3 corresponds to an interval *θ/π* ∈ [0 : 0.34], roughly matching the region of the spectrogram reported in Fig. 11 (g) where the maximum power of slow gamma oscillations is observable. Due to the hysteretic nature of the sub-critical Hopf transition, even if the forcing current *I*_0_(*θ*) decreases for *θ* → 2*π*, we would not observe slow gamma at large *θ*-angles. This is indeed confirmed by the average value of the order parameter *ρ* (grey solid line in Fig. 12 (b)) which reveals an almost monotonic increase with *θ* over a cycle.

For currents *I*_0_ *> I*^(*H*)^ only the limit cycles (corresponding to fast gamma COs with frequencies *ν*_*O*_) arestable, indeed the maximum of the power spectrum for fast gamma COs occurs for *θ* ≃ *π* where *I*_0_ ≃ 0.6 *> I*^(*H*)^ is maximal.

As a last point, let us examine if the coexistence of fast and slow gamma COs is related to some form of P-P locking between theta forcing and gamma oscillations [35, 40]. As evident from Fig. 13 (a) and (b) the theta forcing at *ν*_*θ*_ = 10 Hz locks the collective network dynamics, characterized by the mean membrane potential and by the *γ*-phase defined in Section II C. In particular, for this specific time window we observe for each *θ*-oscillations exactly six *γ*-oscillations of variable duration: slower at the extrema of the *θ*-window and faster in the central part. In agreement with the expected coexistence of *γ* rhythms of different frequencies.

**FIG. 13.**
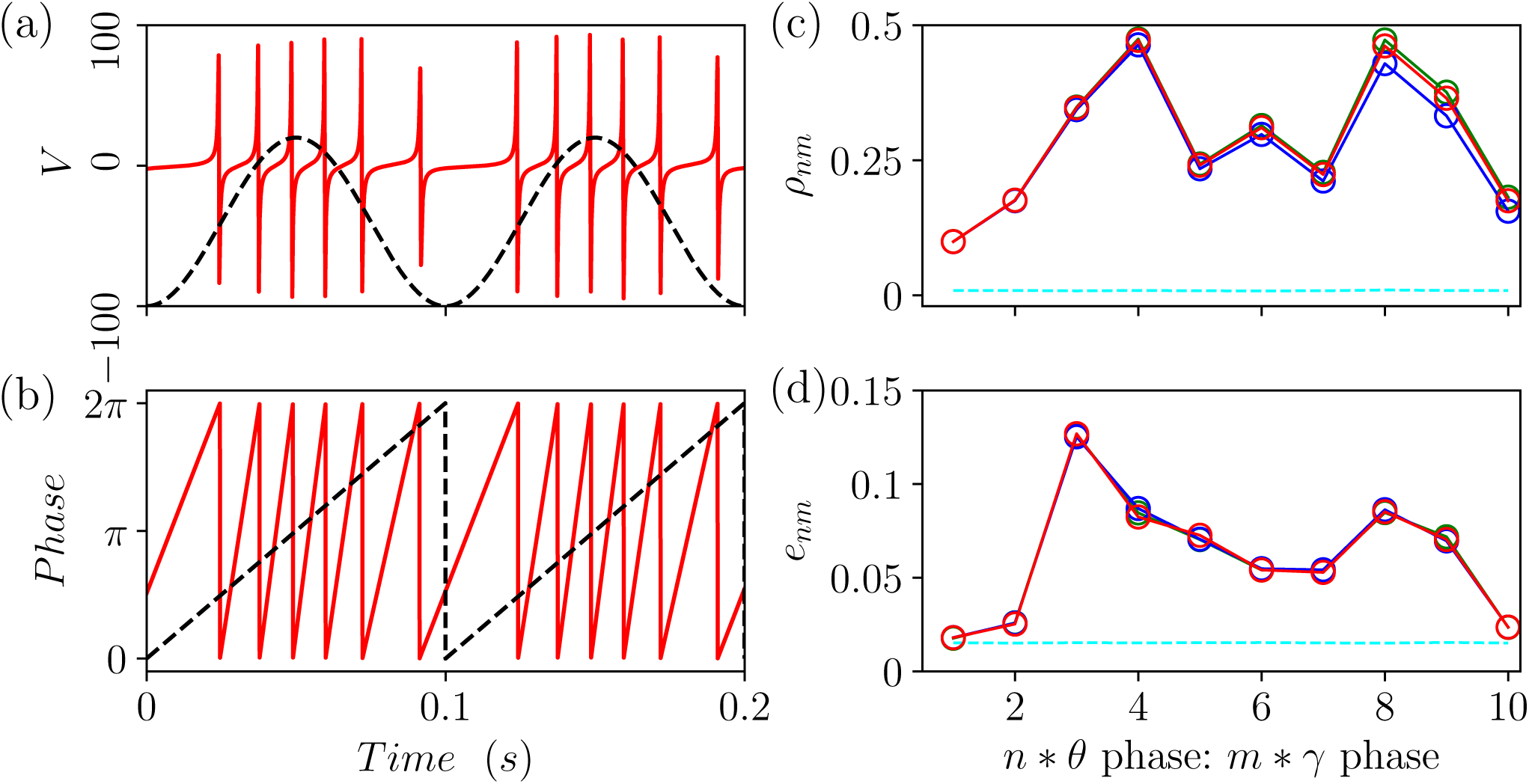
Phase-phase coupling *n* : *m* between theta forcing and gamma oscillations. (a-b) Locking of the gamma oscillations to the external theta forcing: (a) average membrane potentila *V* versus time, the black dashed line is the forcing (13) in arbitrary units; (b) gamma (red solid) and theta (black dashed) phases for the corresponding time interval. (c) Kuramoto order parameter *ρ*_*nm*_ for the phase difference Δ_*nm*_(*t*) for time windows of duration *T*_*W*_ = 0.1 s (black), 0.5 s (red) and 1 s (blue) averaged over 70 *< M <* 700 different realizations. (d) Normalized entropy *e*_*nm*_ for a time window *T*_*W*_ = 0.5 s averaged over *M* = 140 realizations (black), surrogate data are also reported corresponding to random permutation (red) and time shift (blue) of the original data averaged over *M* = 100 independent realizations. The reported data refer to the simulation of the spiking network subject to the external forcing (13) with additive noise on the membrane potentials. Parameters are the same as in Fig. 11, apart for *I*_*θ*_ = 0.3, *ν*_*θ*_ = 10 Hz and *A*_*n*_ = 1.0 × 10^−3^, the histogram of Δ_*nm*_(*t*) employed for the estimation of *e*_*nm*_ have been evaluated over *M* = 50 bins. The results refer only to phases associated to gamma frequencies in the band 30 *–* 100 Hz. The the error bars in (c) and (d) are of the order of the size of the symbols and the significance levels are reported as dashed cyan lines in (b) *ρ*^(*S*)^ = 0.009 and (c) *e*^(*S*)^ = 0.016.

Let us quantify these qualitative observations by considering statistical indicators measuring the level of *n* : *m* synchronization for irregular/noisy data over a large number of theta cycles. In particular, we will employ the Kuramoto order parameter *ρ*_*nm*_ and the normalized entropy *e*_*nm*_ introduced in Section II C measured over time windows of duration *T*_*W*_ and averaged over many different realizations.

As shown in Fig. 13 (c) and (d), both these indicators exhibit two maxima showing the existence of two different locking between *θ* and *γ* oscillations for *n* : *m* equal to ≃ 3 − 4 and ≃ 8, thus corresponding to slow and fast gamma (being *ν*_*θ*_ = 10 Hz). By following [63], in order to test if the reported P-P couplings are significant, we have estimated *ρ*_*nm*_ over time windows of increasing duration, namely from 0.1 s to 1 s. As shown in Fig. 13 (c) the measured values do not vary substantially even by increasing *T*_*W*_ by a factor 10. This is a clear indication of the stationarity of the P-P locking phenomenon here analysed [63]. Furthermore, we measured *e*_*nm*_ also for surrogate data obtained by random permutation and by time-shift (for the exact definitions see Section II C and [63]), the values obtained for these surrogate data are almost indistinguishable from the original ones (see Fig. 13 (d)). These results demand for the development of more effective approaches able to distinguish true locked state from spurious locking.

Finally, the significance level of the reported measurements have been evaluated by randomly shuffling the time stamps of the *γ*-phases and denoted as *ρ*^(*S*)^ and *e*^(*S*)^, respectively (dashed lines in Fig. 13 (c) an (d)). The values of *ρ*^(*S*)^ and *e*^(*S*)^ are definitely smaller than those of the corresponding indicators in correspondence of the observed P-P lockings, thus confirming their significance.

## VI. CONCLUSIONS

In this paper we have shown in terms of an effective mean-field that in a sparse balanced inhibitory network with finite synaptic decay COs can emerge via super or sub-critical Hopf bifurcations from a stable focus. Furthermore, in the network (for sufficiently low structural heterogeneity) the macroscopic focus turns out to be unstable towards microscopic fluctuations in the firing activity leading to the emergence of COs characterized by a frequency corresponding to that of the damped oscillations towards the MF focus. Therefore in proximity of the sub-critical Hopf bifurcations the coexistence of two COs with different origins is observable: slow (fast) gamma oscillations being fluctuation (mean) driven.

From our analysis it emerges that two ingredients are needed to observe coexisting slow and fast gamma COs: the sparsness in the connections and the dynamical balance of the network activity. In particular, the sparsness has a twofold effect at the macroscopic and at the microscopic level. In a mean-field framework the randomness in the in-degree distribution can be reinterpreted as a quenched disorder in the synaptic couplings, which gives rise to the coexistence of stable foci and limit cycles. However, in a fully coupled network with heterogeneous parameters we would not observe strong irregular fluctuations at the level of single neurons, analogous to Poissonian-like firings ususally observed in the cortex [56, 57, 81]. These can emerge only in sparsely connected networks [19, 20]. Moreover, the balance mechanism guarantees that the irregular spiking dynamics will not disappear in the thermodynamic limit [21, 23–25].

These persistent microscopic fluctuations are able to trigger slow gamma COs in the network, which coexist with fast gamma COs corresponding to the limit cycle solutions in the MF. These two ingredients usually characterize real brain networks, where our prediction that slow (fast) gamma oscillations are associated to more (less) irregular neuronal dynamics can be experimentally tested e.g. by measuring the coefficient of variation associated to these two states. Furthermore, previous theoretical analysis of gamma oscillations based on two interacting Wilson-Cowan rate models with different synaptic times revealed only the possible coexistence of two stable limit cycles both corresponding to tonic collective firing (i.e. mean driven COs) [54].

A fundamental improvement of the effective mean-field here introduced should include in the formulation also the current fluctuations due to the sparsness. A possible strategy to incorporate these intrinsic noise sources in an exact mean-field formulation, going beyond the Ott-Antonsen Ansatz [8], could rely on the circulant cumulants expansion recently applied to ensemble of oscillators [82] and of QIF neurons [83] in presence of extrinsic noise.

Furthermore, a standard approach to obtain a macroscopic description of the neuronal network dynamics is based on derivation of the Fokker-Planck equation for the distribution of the membrane potentials. This formalism has been fully developed for sparse networks of Leaky Integrate-and-Fire neurons in [19, 20], preliminary analysis based on this approach for the QIF network examined here and in [21] suggest that fluctuation driven COs emerge due to an instability of the asynchronous state [84].

Our model is not meant to explicitly replicate the dynamics of specific brain areas, but rather to illustrate fundamental mechanisms by which slow and fast gamma oscillations may arise and coexist due to local network inhibitory features. However, several phenomena we reported resemble experimental results obtained for different brain regions *in vitro* as well as *in vivo* and our findings can stimulate new experiments or lead to novel interpretation of the existing data.

Of particular interest is the possibility, analysed in Section V A, to switch from a gamma rhythm to the other by performing transient stimulations. This mechanism can allow a single inhibitory population to pass from a coding task to another following an external sensory stimulus. Indeed it has been shown that distinct gamma rhythms are involved in different coding processes: namely, fast gamma in new memory encoding, while slow gamma has been hypothized to promote memory retrieval [85].

On one side, pathological synchronization is usually associated to neuronal diseases [15, 86, 87]. On another side, aberrant gamma oscillations have been observed in several cognitive disorders, including Alzheimer’s disease, Fragile X syndrome and neocortical epilepsy [79, 85]. Furthermore, deep brain stimulation (DBS) techniques have been developed along the years to treat some of these diseases, e.g. essential tremor and Parkinsons’ disease [88–90]. We have presented a simple model exhibiting the coexistence of highly synchronized states and asynchronous or partially synchronized regimes. Therefore, our model can represent a simple benchmark where to test new DBS protocols to obtain eventually less invasive technique to desynchronize pathological states [91–93] or to restore healthy gamma rhythms, as suggested in [85].

Moreover, the richness of the dynamical scenario present in this simple model indicates possible future directions where intrinsic mechanisms present in real neural networks like spike frequency adaptation could permit a dynamical alternation between different states. In this direction, a slow variable like adaptation could drive the system from “healthy” asynchronous or oscillatory dynamics to periods of pathological extremely synchronous regimes, somehow similar to epileptic seizure dynamics [94].

In Section V B, we have analysed the emergence of COs in our network in presence of an external theta forcing. This in order to to make a closer contact with recent experimental investigations devoted to analyse the emergence of gamma oscillations in several brain areas *in vitro* under sinusoidally modulated theta-frequency optogenetic stimulations [39, 48, 49]. For low forcing amplitudes, our network model displays theta-nested gamma COs at frequencies ≃ 50 Hz joined with irregular spiking dynamics. These results are analogous to the ones reported for the CA1 and CA3 areas of the hippocampus in [39, 48], moreover theta-nested oscillations with similar features have been reported also for the mEC [49], but for higher gamma frequencies.

Furthermore, for intermediate forcing amplitudes we observe the coexistence of slow and fast gamma oscillations, which lock to different phases of the theta rhythm, analogously to what reported for the rat hippocampus during exploration and REM sleep [35, 46]. The theta-phases preferences displayed in our model by the different gamma rhythms are due to the hysteretic nature of the sub-critical Hopf bifurcation crossed during the theta stimulation. Finally, for sufficiently strong forcing, the model is driven in the fast gamma regime.

Our analysis suggests that a single inhibitory population can generate locally different gamma rhythms and lock to one or the other in presence of a theta forcing. In particular, we have shown that fast gamma oscillations are locked to a strong excitatory input, while slow gamma COs emerge when excitation and inhibition balance. These results can be useful in revealing the mechanism behind slow and fast gamma oscillations reported in several brain areas: namely, hippocampus [45], olfactory bulb [95], ventral striatum [96], visual cortical areas [97] and neocortex [44]. Particularly interesting are the clear evidences reported in [44] that different gamma rhythms, phase locked to the hippocampal theta rhythm, can be locally generated in the neocortex. Therefore future studies could focus on this brain region to test for the validity of the mechanisms here reported.

For what concerns the CA1 area of the hippocampus, where most of the experimental studies on theta-gamma oscillations have been performed. Despite the experimental evidences that different gamma oscillations emerging in CA1 area at different theta phases are a reflection of synaptic inputs originating from CA3 area and mEC [46, 47] this does not exclude the possibility that a single CA1 inhibitory population can give rise to different gamma rhythms depending on the network state [45]. This hypothese is supported by experimental evidences showing that a large part of CA1 interneurons *in vivo* can lock to both slow and fast gamma [35, 46, 47] and that *in vitro* gamma rhythms can be locally generated in various regions of the hippocampus due to optogenetic stimulations [39, 48, 49] or pharmalogical manipulation [50–53]. However, much work remains to be done to clarify if local mechanisms can give rise to coexisting gamma rhythms also in the CA1 area.

Another interesting aspect, revealed quite recently, concerns the wide cycle-to-cycle variability in the theta-nested gamma oscillations observed during spatial exploration and memory-guided behaviour in CA1 [80] and in dentate gyrus [98]. In particular, in these papers the authors show not only that different gamma bands can be excited during different theta cycles, but also that the spectral range of these bands overlap. This is consistent with our analysis for intermediate forcing amplitude reported in Figs. 11 and 12, where on one side we show that the dynamics from one cycle to the next one is extremely variable and on another side that the frequency bands associated to slow and fast COs slightly overlap around 40 Hz.

At variance with previous results for purely inhibitory populations reporting noise sustained COs in the range 100–200 Hz [16] our model displays slow gamma rhythms characterized by irregular firing of the single neurons. Therefore in our case it is not necessary to add an excitatory population to the inhibitory one to slow down the rhythm and to obtain oscillations in the gamma range as done in [30, 99]. Evidences have been recently reported pointing out that gamma oscillations can emerge locally in the CA1 induced by the application of kainate due to purely inhibitory mechanisms [53]. However, other studies point out that in the same area of the hippocampus excitatory and inhibitory neurons should interact to give rise to oscillations in the gamma range [39, 52]. Preliminary results obtained for QIF networks with a sinusoidal theta forcing show that theta-nested gamma oscillations with similar features can emerge for purely inhibitory as well as for mixed excitatory-inhibitory networks [100].

As shown in Section III B the same kind of bifurcation diagram can be observed by considering the external excitatory drive as well as the self-disinhibition of the recurrently coupled inhibitory population. This suggests that in our model the same scenarios reported in Section V for an excitatory theta forcing can be obtained by considering an external inhibitory population which transmits rhythmically its activity to the target population. This somehow mimicks the pacemaker theta activity of a part of the medial septum interneurons on the interneurons of the hippocampus experimentally observed in [101]. This subject will be addressed in future studies due to its relevance in order to clarify the origin of theta-gamma oscillations in the hippocampus, however it goes beyond the scopes of the present analysis.

In this paper we considered a model including the minimal ingredients necessary to reproduce the phenomenon of coexisting gamma oscillations corresponding to quite simple (namely, periodic) collective regimes. However, the introduction of synaptic delay in the model can lead to more complex coexisting states, like quasi-periodic and even chaotic solutions, as recently shown for fully coupled networks in [65, 102]. The inclusion of delay in our model can enrich the dynamical scenario maybe allowing to mimic further aspects of the complex patterns of activity observed in the brain, like e.g. sharp-wave ripples observed in the hippocampus and which are fundamental for memory consolidation [103]. Due to the large variety of interneurons present in the brain and in particular in the hippocampus [104] a further step in rendering our model more realistic would consist in considering multiple inhibitory populations characterized by different neuronal parameters. By manipulating the influence of a population on the others it would be interesting to investigate the possible mechanisms to switch COs from one gamma rhythm to another, following the richness of the bifurcation scenarios presented in Figs. 2 and 3.

The generality of the phenomena here reported is addressed in Appendix A and B. In particular in Appendix A we show that the mechanisms leading to the coexistence scenario of fast and slow gamma oscillations are not peculiar of Lorentzian in-degree distributions (that we employed to allow a comparison of the network simulations with the MF results), but that they are observable also in the more studied Erdös-Reniy sparse networks. Appendix B is devoted to the analysis of a suitable normal form, which reproduces the dynamics of the MF in proximity of the sub-critical Hopf bifurcation. In particular, the noisy dynamics of the normal form reveals coexisting oscillations of different frequencies. More specifically the addition of noise leads from damped oscillations towards the stable focus to sustained oscillations characterized by the same frequency. This latter result links our findings to the more general context of noise-induce oscillations for non-excitable systems examined in various fields of research: namely, single cell oscillations [105], epidemics [106], predator-prey interactions [107] and laser dynamics [108]. At variance with all previous studies we have analyzed noise-induced oscillations coexisting and interacting with oscillations emerging from the Hopf bifurcation. Furthermore, the mechanism leading to the irregular fluctuations in our case is quite peculiar. Single cells oscillations are believed to be driven by molecular noise, induced by the small number of molecules present in each cell, and therefore disappearing in the thermodynamic limit [109]. Recently, another possible mechanism leading to fluctuation amplification in a feed-forward chain has been suggested as a pacemaking mechanism for biological systems, in this context the amplitude of the oscillations grows with the system size [110]. Instead in our case, the dynamical balance provides intrinsic noise and oscillations of constant amplitude, essentially independent from the number of synaptic inputs (in-degree) and from the number of neurons in the network.

## ACKNOWLEDGMENTS

We acknowledge useful discussions with D. Avitabile, D. Angulo-Garcia, D. Battaglia, F. Devalle, S. Keeley, E. Montbrió, S. Olmi, A. Politi, J. Rinzel, and R. Schmidt. The authors received partial economic support by the French Governement via the Excellence Initiative I-Site Paris Seine (Grant No ANR-16-IDEX-008), the Labex MME-DII (Grant No ANR-11-LBX-0023-01) and the ANR Project ERMUNDY (Grant No 18-CE37-0014-03).

## Appendix A Slow and fast gamma oscillations in Erdös-Reniy network

In order to compare the network simulations with the MF results we have considered in the article a Lorentzian distribution for the in-degrees. It is therefore important to show that the same phenomenology is observable by considering a more standard distribution, like the Erdös-Reniy (ER) one. The results of adiabatic simulations, reported in Fig. 14, confirm that also for ER networks a bistable regime, characterized by COs with different gamma-frequencies (see panel (b)), is indeed observable. In particular, slow gamma COs characterized by an average firing rate 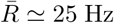 and irregular neuronal firings (as shown in panels (c) and (e)) coexist with almost synchronized fast gamma COs with neurons tonically firing with 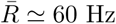 (see panels (d) and (f)).

**FIG. 14.**
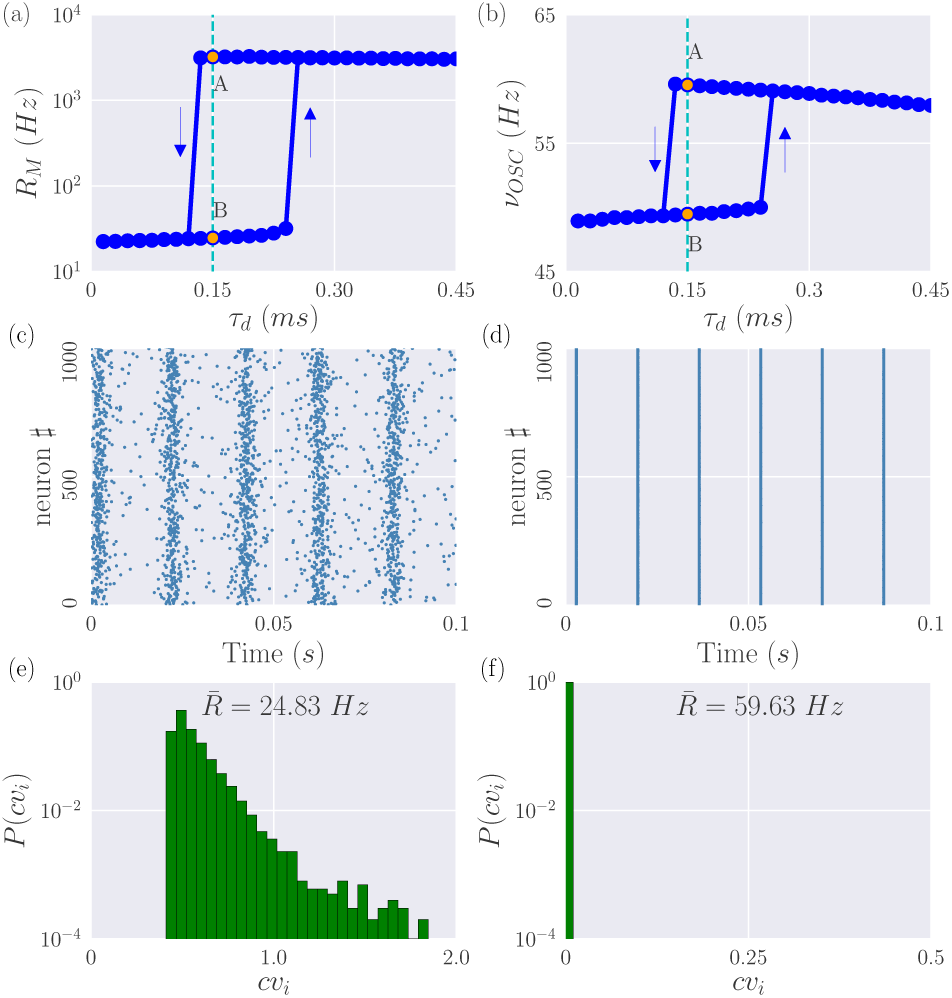
Erdös-Reniy Network. Results of adiabatic simulations for an ER network obtained by varying the synaptic time *τ*_*d*_: (a) maximal firing rates *R*_*M*_ and (b) frequencies *ν*_*OSC*_ of the COs. Two coexisting states (A) and (B) are considered at *τ*_*d*_ = 0.15 ms. In the left and right row are reported the raster plots (c,d) and the distributions of the *cv*_*i*_ (e,f) for the state (A) and (B), respectively. Parameters for the simulations are *N* = 10000, *K* = 1000, *I*_0_ = 0.25, *J*_0_ = 1.0 and Δ*τ*_*d*_ = 0.015 ms, 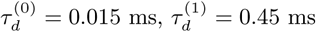.

## Appendix B A general mechanism for the emergence of coexisting oscillations

We investigate here the generality of the mechanism for the coexistence of COs observed in the network of QIF neurons. In particular, we have shown that this phenomenon occurs when in the MF model we have a focus coexisting with a limit cycle, while in the sparse network we have fluctuations sustained by the dynamical balance. If this is the mechanism we expect to see a similar phenomenon whenever we consider a system in proximity of a sub-critical Hopf bifurcation and we add noise of constant amplitude to the dynamics.

Therefore, to asses the generality of the phenomenon we consider the normal form of a Hopf bifurcation in two dimensions leading to the birth of a limit cycle from an equilibrium, namely [111, 112]:

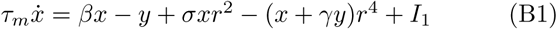

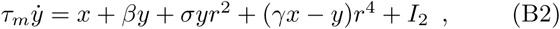

where *r*^2^ = *x*^2^ + *y*^2^, *τ*_*m*_ = 4 ms is an arbitrary time scale, *I*_1_(*t*) and *I*_2_(*t*) are generic external time dependent forcing, *β* is the bifurcation parameter, the parameter *σ* sets the nature of the bifurcation and *γ* controls the frequency of the stable and unstable limit cycles. Notice that we added a quintic term, absent in the original normal form [111, 112], in order to maintain bounded the values of *x* and *y* while keeping the same bifurcation structure. For *I*_1_ = *I*_2_ = 0 we will have a sub-critical (super-critical) Hopf for *σ* = +1 (*σ* = −1). In this case it is convenient to rewrite (B2) in polar coordinates (*x, y*) = (*r* cos *ϕ, r* sin *ϕ*), as follows:

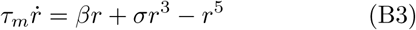

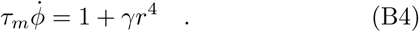

The stationary solutions are *r* = 0 corresponding to stable focus characterized by relaxation oscillations with a frequency *ν*_*D*_ ≃ 39 Hz and a stable and unstable limit cycles of amplitudes 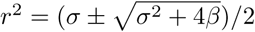.

In Fig. 15 (a) we report the bifurcation diagram for *σ* = +1 and *I*_1_ = *I*_2_ = 0. We observe that the subcritical Hopf bifurcation occurs at *β* = *β*_*c*_ = 0 and for *β <* 0 it exists a region where a stable (green dots) and unstable (blue dashed line) limit cycles coexists with a stable focus (red line), exactly as it happens for the QIF MF model (see Fig. 7 (a)). The stable and unstable limit cycles merge at a SN bifurcation located at *β* = −*σ*^2^/4.

**FIG. 15.**
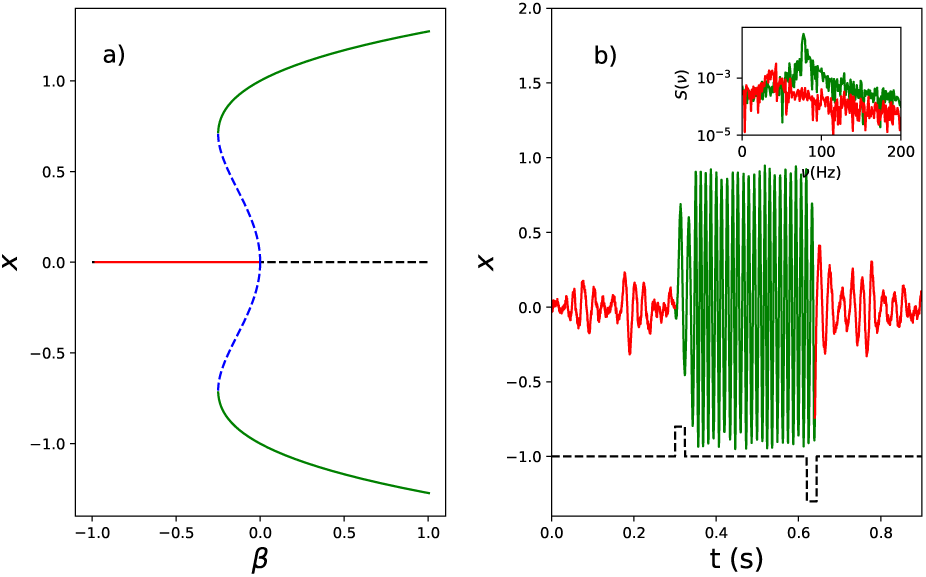
a) Bifurcation diagram for the variable *x* as a function of the parameter *β*. Green (blue) lines indicate a stable (unstable) limit cycle and red (black) line a stable (unstable) focus. b) Fixing *β* in the bistability region we report the time trace of *x*(*t*) in presence of a zero-mean gaussian noise of amplitude *A*_1_ = *A*_2_ = 0.14. An external pulse of current is added to the evolution equation for *x* in (B2) for a time window of 56 ms to induce a switching between the oscillatory states (the black dashed line, shifted on the *x* axe to be visible while the actual baseline value is zero). In the inset we report the power spectrum of the two different oscillatory regimes obtained over long time traces (hundreds of seconds) in order to check that the oscillations persist in time. Parameters are *β* = −0.16, *σ* = 1, *γ* = 1.5.

As previously stated, the MF model cannot capture the endogenous fluctuations, naturally present in sparse balanced networks. In order to emulate this effect we consider *I*_1_(*t*) and *I*_2_(*t*) to be two i.i.d. Gaussian white noise processes (i.e. *I*_*q*_(*t*) = *A*_*q*_*ξ*_*q*_(*t*) with *q* = 1, 2, where *ξ*_*q*_(*t*) are random, Gaussian distributed, variables of zero average and unitary variance). In presence of these additive noise terms and in proximity of the Hopf bifurcation, we observe the coexistence of two oscillatory regimes as shown in Fig. 15 (b). One oscillation, characterized by higher amplitude (green line), corresponds to the limit cycle present in the non-noisy dynamics (green line in the bifurcation diagram reported in Fig. 15 (b).). The other oscillation is the result of a constructive role of noise that excites the stable focus thus generating robust oscillations at the frequency *ν*_*D*_ (red line). Analogously to what shown for the network of QIF neurons (see Fig. 8), it is possible to switch between the two kind of oscillations via a pulse current of positive (negative) amplitude with respect to the baseline (see the dashed line in panel b)). Moreover the frequencies of the two oscillations, generated by two different mechanisms, corresponds to slow and fast gamma oscillations as observable in the corresponding power spectra *S*(*ν*) reported in the inset of Fig. 15 (b)..

